# Eosinophils protect against SARS-CoV-2 following a vaccine breakthrough infection

**DOI:** 10.1101/2024.08.08.607190

**Authors:** Kathryn M. Moore, Stephanie L. Foster, Meenakshi Kar, Katharine A. Floyd, Elizabeth J. Elrod, M. Elliott Williams, Jacob Vander Velden, Madison Ellis, Ansa Malik, Bushra Wali, Stacey Lapp, Amanda Metz, Steven E. Bosinger, Vineet D. Menachery, Robert A. Seder, Rama Rao Amara, Jacob E. Kohlmeier, Arash Grakoui, Mehul S. Suthar

**Affiliations:** Emory Vaccine Center, Emory National Primate Research Center, Emory University, Atlanta, Georgia, USA; Center for Childhood Infections and Vaccines of Children’s Healthcare of Atlanta, Department of Pediatrics, Emory University School of Medicine, Atlanta, Georgia, USA; Emory Center of Excellence of Influenza Research and Response (CEIRR), Atlanta, Georgia, USA; Department of Microbiology and Immunology, Emory University, Atlanta, Georgia, USA; Vaccine Research Center, National Institute of Allergy and Infectious Diseases, National Institutes of Health, Bethesda, Maryland, USA; Emory NPRC Genomics Core Laboratory, Atlanta, GA; Department of Medicine, Emory University School of Medicine, Atlanta, Georgia, USA

**Author notes:** Correspondence: Mehul S. Suthar.

## Abstract

Waning immunity and the emergence of immune evasive SARS-CoV-2 variants jeopardize vaccine efficacy leading to breakthrough infections. We have previously shown that innate immune cells play a critical role in controlling SARS-CoV-2. To investigate the innate immune response during breakthrough infections, we modeled breakthrough infections by challenging low-dose vaccinated mice with a vaccine-mismatched SARS-CoV-2 Beta variant. We found that low-dose vaccinated infected mice had a 2-log reduction in lung viral burden, but increased immune cell infiltration in the lung parenchyma, characterized by monocytes, monocyte-derived macrophages, and eosinophils. Single cell RNA-seq revealed viral RNA was highly associated with eosinophils that corresponded to a unique IFN-γ biased signature. Antibody-mediated depletion of eosinophils in vaccinated mice resulted in increased virus replication and dissemination in the lungs, demonstrating that eosinophils in the lungs are protective during SARS-CoV-2 breakthrough infections. These results highlight the critical role for the innate immune response in vaccine mediated protection against SARS-CoV-2.

## Introduction

In response to the COVID-19 pandemic, vaccines against severe acute respiratory syndrome coronavirus 2 (SARS-CoV-2) were rapidly developed and demonstrated protection from severe disease, hospitalization, and death from COVID-19. Neutralizing antibodies induced by vaccination are an immune correlate of protection against COVID-19 as higher serum neutralizing antibody titers are associated with protection from symptomatic disease (*1, 2*). However, COVID-19 vaccine-induced antibodies wane over the course of six months which results in reduced vaccine effectiveness (*3–5*). When neutralizing antibody titers wane, COVID-19 vaccination provides protection from infection in the lower respiratory tract but do not provide protection from infection in the upper respiratory tract (*2, 6, 7*). Furthermore, SARS-CoV-2 continues to evolve leading to the emergence of genetically divergent variants from the ancestral virus. As a result, serum antibodies induced by COVID-19 vaccination and previous infections have reduced neutralizing activity against variants compared to ancestral SARS-CoV-2 (*8–11*). Reduction in neutralization due to waning antibody titers over time and the emergence of immune evasive variants allows SARS-CoV-2 to establish infection, which we define as a breakthrough infection (*1, 12, 13*).

The innate immune system is responsible for sensing and responding to pathogens and coordinating the adaptive immune response (*14*). Early studies during the pandemic of unvaccinated COVID-19 patients revealed that innate immune cells including monocytes, neutrophils, and dendritic cells infiltrate the lungs during SARS-CoV-2 infection (*15–17*). This has also been recapitulated in mouse models of SARS-CoV-2 (*18–22*). We have previously shown that monocytes are an essential component of the immune response to SARS-CoV-2, where loss of critical monocyte signaling resulted in elevated viral load in the lungs and 60% mortality in mice infected with mouse-adapted SARS-CoV-2 (*19*). In experimental models, neutrophil extracellular traps have been shown to mediate lung epithelial cell damage and neutrophil proteases have been shown to degrade SARS-CoV-2 spike protein and abrogate virus replication, demonstrating that neutrophils may contribute to both pathogenicity and protection against SARS-CoV-2 (*23–25*). However, the innate immune response following a SARS-CoV-2 vaccine breakthrough infection has not yet been well characterized.

To model breakthrough infections in mice, we vaccinated mice with a low-dose of vaccine against the SARS-CoV-2 spike protein based on the ancestral Wuhan-1 strain followed by vaccine-mismatched challenge with the Beta variant (B.1.351). We vaccinated 129S1/SvImJ mice with either prefusion-stabilized spike protein subunit vaccine adjanted with Addavax or mRNA vaccine encoding prefusion-stabilized spike protein. Using flow cytometry and single cell RNA sequencing (scRNA-seq), we observed a distinct innate immune response to breakthrough infection in the lungs compared to naïve (unvaccinated) infected mice. We observed a higher level of innate immune cells in the lungs during SARS-CoV-2 breakthrough infections compared to naïve infected mice but with reduced activation marker expression. Eosinophils infiltrated the lungs at a higher frequency in vaccinated mice as compared to naïve mice, regardless of vaccine platform. Depleting eosinophils in vaccinated mice resulted in reduced protection of the lungs after B.1.351 infection. These results indicate a critical role for innate immunity, triggered by vaccination, in viral control. Furthermore, these results demonstrate that eosinophils play a protective role during SARS-CoV-2 infection.

## Results

### S-2P protein subunit vaccination of mice protects against SARS-CoV-2 infection

Mice were vaccinated with a prefusion-stabilized spike protein based on the Wuhan-1 strain (hereafter referred to as S-2P) to reflect the neutralizing antibody titers against early B lineage SARS-CoV-2 variants generally observed in humans following COVID-19 vaccination (*26*). Protein subunit vaccines comprising prefusion-stabilized SARS-CoV-2 spike protein combined with aluminum hydroxide adjuvant were found to be efficacious in preclinical testing in mice and clinical trials in humans (*27, 28*). Furthermore, the S-2P antigen has been licensced for use as a subunit vaccine in Taiwan (MCV-COV1901) (*29*). To model waning immunity in mice, we vaccinated mice with either 5 or 0.5 micrograms (μg) of S-2P adjuvanted with AddaVax, an MF59-like adjuvant, via the subcutaneous route, and each mouse received a boost of the same formulation 30 days later. At 30 days post-boost, S-2P-vaccinated mice and naïve age-matched controls were infected with 1 x 10^6^ plaque-forming units (PFU) of B.1.351 through the intranasal (i.n.) route. A schematic of the vaccine schedule and challenge are shown in **Figure 1A**.

**Figure 1.**
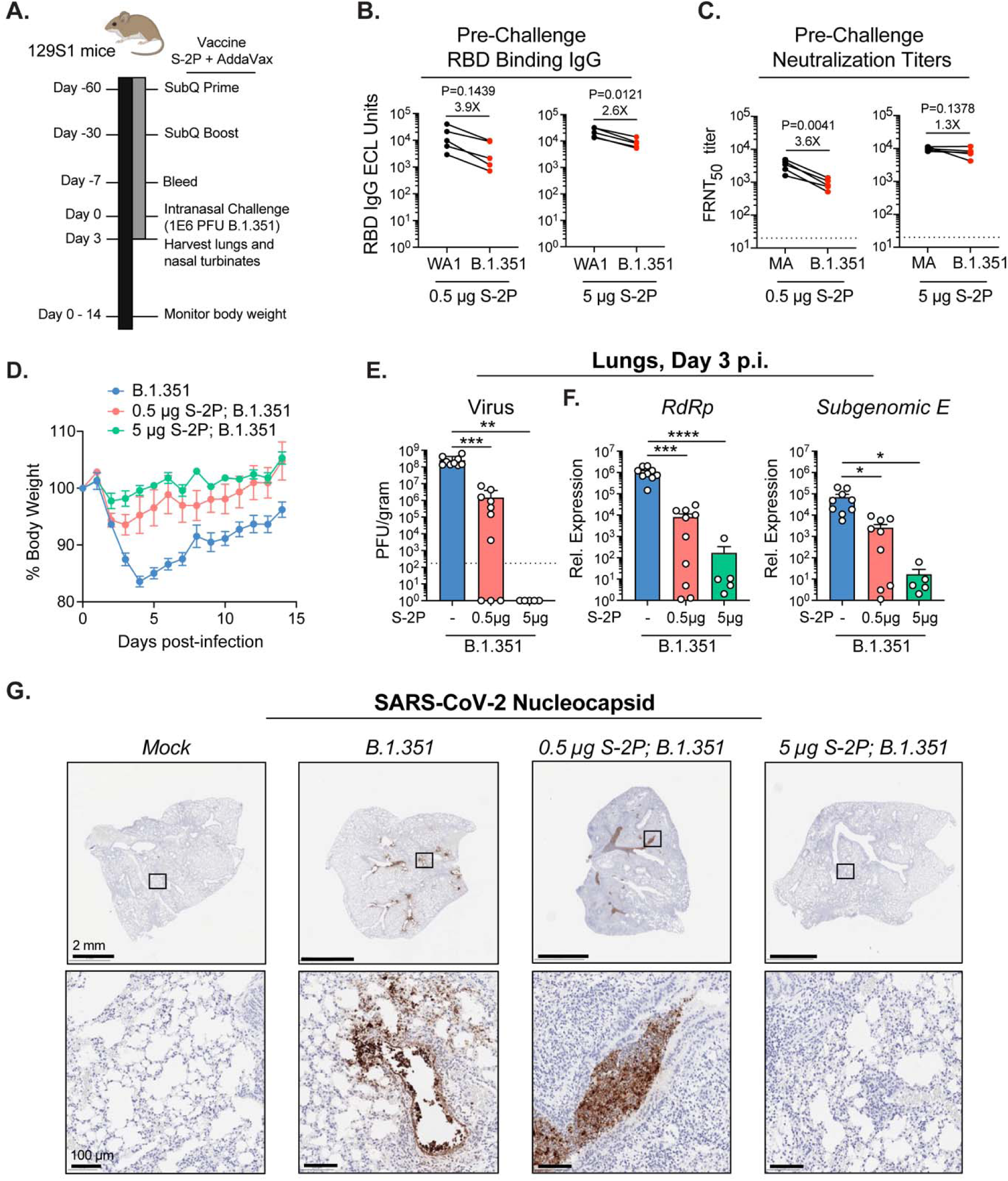
Protein subunit vaccination of mice protects against SARS-CoV-2 infection. (**A**) Experimental schematic detailing vaccination of 129S1/SvImJ mice with S-2P and subsequent challenge with B.1.351. (**B**) Live virus serum FRNT_50_ titers against MA-SARS-CoV-2 (black circles) or B.1.351 (red circles) at 7 days before challenge. Connected circles represent the same mouse. The dotted horizontal line represents the LOD of the assay. (**C**) Serum IgG binding antibodies specific for SARS-CoV-2 RBD **(C)** measured 7 days before challenge using a Meso Scale Discovery V-Plex SARS-CoV-2 kit. The ECL units for IgG binding to WA-1 (black circles) and B.1.351 (red circles) RBD from individual mice are connected by a line. The P value and fold difference in geometric mean titer (GMT) between WA-1 or MA-SARS-CoV-2 and B.1.351 are shown for (**B-C**). (**D**) Body weight loss as a percentage of day of challenge for mice infected with B.1.351 following vaccination with 0.5 µg S-2P, 5 µg S-2P, or naïve age-matched controls. Symbols represent the mean percent body weight loss of five mice. (**E-F**) B.1.351 viral loads in the lungs as measured by plaque assay (**E**) and relative expression of genomic (RdRp) and subgenomic (E) viral RNA determined by RT-qPCR (**F**) at day 3 p.i. The dotted horizontal line represents the LOD of the assay. (**G**) Histopathological analysis of mouse lungs harvested at day 3 p.i. stained with an antibody specific to SARS-CoV-2 nucleoprotein (N). Representative images of SARS-CoV-2 N staining. Scale bars are 2 mm (top) and 100 µm (bottom). Group names, sample size, and color are as follows: uninfected/mock, n=5, grey; naïve infected/B.1.351, n=4, blue; low-dose vaccinated infected/0.5 µg S-2P, B.1.351, n=5, red; high-dose vaccinated infected/5 µg S-2P, B.1.351 = green. Data are represented by the mean +/- the standard error of the mean. Statistical significance was determined using an unpaired Student’s *t* test, and P values are represented as follows: **, P < 0.01; ***, P < 0.001. Created with BioRender.com.

S-2P-vaccinated mice developed binding antibodies to the spike receptor-binding domains (RBDs) of the closely vaccine-matched WA-1 variant and of the challenge variant B.1.351 and neutralizing antibodies (**Figure 1B-C**). Binding antibody titers in 0.5 μg S-2P S-2P-vaccinated mice were modestly lower (3.9-fold; P = 0.1439) for B.1.351 than WA-1, reflecting the sequence divergence of B.1.351 from the Wuhan-1 spike used in the protein subunit vaccine, as previously reported by our group (*10*) (**Figure 1B**). The mouse-adapted (MA) SARS-CoV-2 strain CMA3p20 (hereafter referred to as MA-SARS-CoV-2) was used to test neutralization of vaccinated mouse sera and produces similar neutralization titers to WA-1 (*30*). We have previously shown that the spike mutations within CMA3p20 does not impact neutralization of ancestral spike vaccine sera as compared to WA-1 (*30*). Vaccinated mice developed neutralizing antibodies against MA-SARS-CoV-2 and B.1.351 prior to infection, although neutralizing titers to B.1.351 were 3.6-fold lower (P = 0.0041) than to MA-SARS-CoV-2 in the 0.5 μg S-2P-vaccinated mice (**Figure 1C**).

Viral control in the lungs of S-2P-vaccinated infected mice was associated with reduced morbidity in this model compared to naïve infected mice, with a vaccine dose-dependent reduction in weight loss and reduced viral load in the lungs. S-2P-vaccinated mice exhibited reduced weight loss and recovered more quickly after B.1.351 challenge than naïve mice (**Figure 1D**). We observed that 0.5 μg S-2P-vaccinated mice had 172-fold lower viral loads in the lungs at day 3 p.i. than naïve infected mice (**Figure 1E-F**), although no significant difference in viral load was observed between naïve infected and 0.5 μg S-2P vaccination in the nasal turbinates (**Supplemental Figure 1**). Further, 0.5 μg S-2P vaccination represented an intermediate viral load in the lungs between naïve and 5 μg S-2P vaccination which resulted in no detectable infectious virus at day 3 p.i. (**Figure 1E**). Histopathological analysis revealed that SARS-CoV-2 nucelocapsid (N) antigen was found predominately in the bronchi (large airways) in 0.5 μg S-2P-vaccinated mice as compared to naïve infected mice where viral antigen was found in both the bronchi and the alveoli (small airways) (**Figure 1G**). Viral antigen was observed in the bronchial lumen debris in 0.5 μg S-2P-vaccinated lungs further indicates possible engulfment and clearance of the virus by phagocytic cells. There was little to no detectable viral antigen in the lungs of 5 μg S-2P-vaccinated mice at 3 days p.i. Taken together, these results demonstrate that low-dose S-2P vaccination (0.5 μg) is a suitable model that shows virus replication in the lungs in the presence of vaccine-induced immune responses.

### Low-dose S-2P-vaccinated mice exhibit an influx of innate immune cells into the lung following B.1.351 infection

We next evaluated the populations of innate immune cells present in the lung to understand the early immune responses to breakthrough infection after vaccination. Immune cell infiltration into the lungs of naïve and 0.5 μg S-2P-vaccinated mice were observed by immunological staining with hematoxylin and eosin (H&E) at 3 days p.i. (**Figure 2A**). Mice vaccinated with 0.5 μg S-2P had a significantly greater infiltration of immune cells into the parenchyma, cellular debris present in the lumen of the bronchi tissue, and in some cases loss of normal alveolar architecture compared to naïve mice. Perivascular inflammation, bronchial/bronchiolar alveolar degeneration/necrosis, bronchial/bronchiolar inflammation, and alveolar inflammation were each scored on a scale from 1 to 5 for each mouse. The sum of these four pathology scores out of 20 is shown in **Figure 2B**. 0.5 μg S-2P-vaccinated infected mice had significantly higher total pathology scores than mock or naïve infected mice. Lungs from naïve infected mice all had some level of inflammation, as indicated by immune cell infiltration, alveolitis, and/or pneumonitis, in contrast to lungs from uninfected mice, which had little or no discernible inflammation.

**Figure 2.**
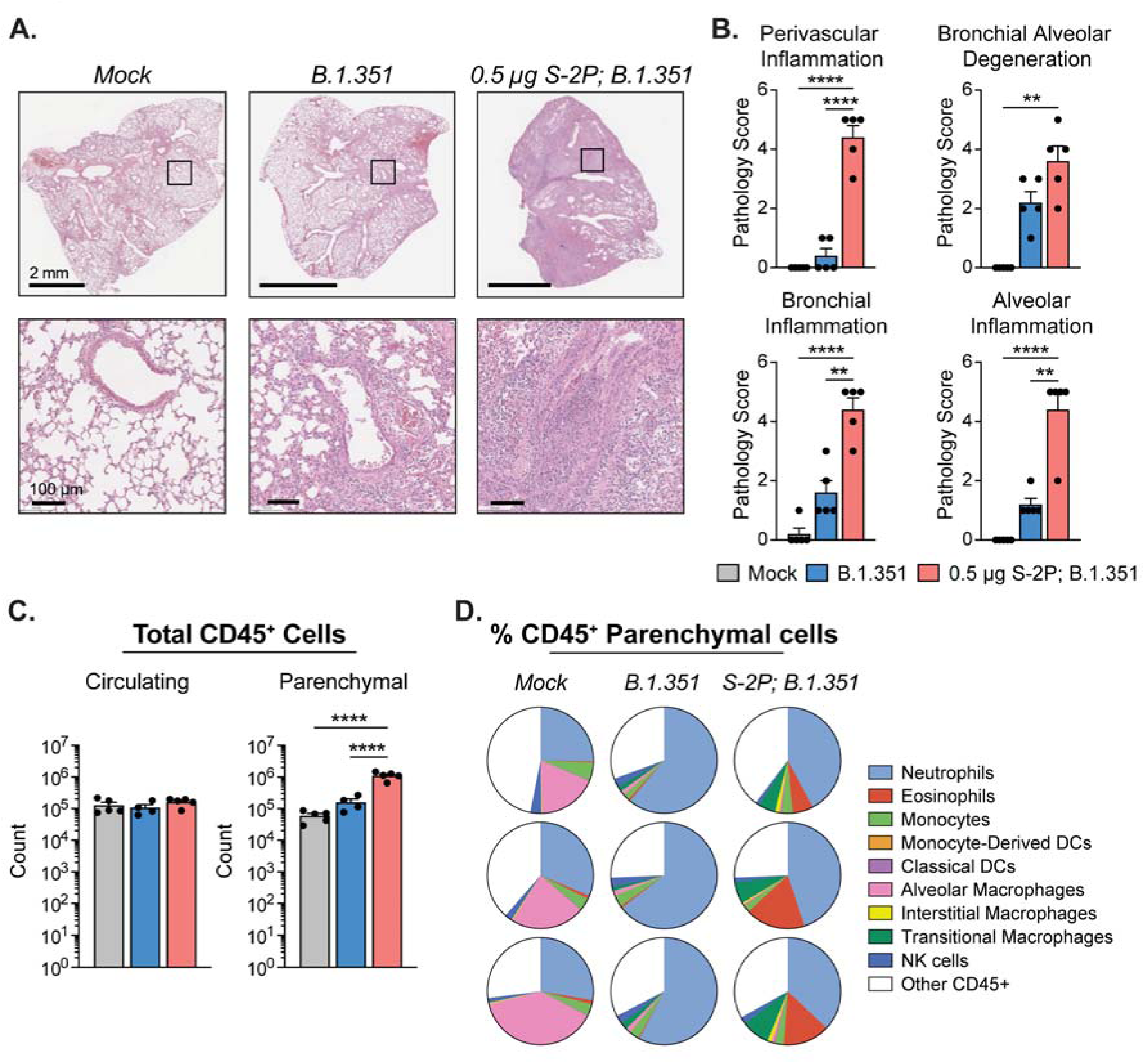
Low-dose S-2P-vaccinated mice exhibit an influx of innate immune cells into the lung following B.1.351 infection. (**A-B**) Histophathologic analysis of mouse lungs harvested at day 3 p.i. stained with H&E. (**A**) Representative images of H&E staining. Scale bars are 2 mm (top) and 100 µm (bottom). (**B**) Bar graphs displaying pathology scores in the following categories: perivascular inflammation, bronchial alveolar degeneration, bronchial inflammation, and alveolar inflammation. (**C**) Absolute counts of CD45^+^ circulating cells (left) and absolute counts of CD45^+^ lung-resident parenchymal cells (right) determined by flow cytometry at day 3 p.i. (**D**) Pie charts displaying proportion of each identified cell type represented as a percent of the total CD45^+^ lung parenchymal cells for 3 representative mice from each group. Group names, sample size, and color are as follows: uninfected/mock, n=5, grey; naïve infected/B.1.351, n=4, blue; low-dose vaccinated infected/0.5 µg S-2P, B.1.351, n=5, red. Data are represented by the mean +/- the standard error of the mean. Statistical significance was determined using an unpaired one-way ANOVA with Tukey’s multiple comparisons test, and P values are represented as follows: ****, P < 0.0001.

We next performed intravital (IV) labeling with an antibody against CD45 conjugated to phycoerythrin (PE) to tag antibody-accessible cells in the lung to distinguish between circulating (PE:CD45^+^) and lung parenchymal (PE:CD45^-^) cells (*19, 31, 32*). We used a multi-parameter flow cytometry panel to define parenchymal CD45^+^ cells (excluding B and T cells) and identify populations of granulocytes (neutrophils and eosinophils), natural killer (NK) cells, monocytes, macrophages, monocyte-derived dendritic cells (moDCs), and classical dendritic cells (cDCs) (**Supplemental Figure 2**). Mock, naïve infected, and 0.5 μg S-2P-vaccinated infected had similar numbers of circulating (CD45 IV-positive) cells in the lungs at day 3 p.i. (**Figure 2C**). However, 0.5 μg S-2P-vaccinated infected mice had 10-fold higher CD45^+^ cells infiltrating into the lung parenchyma as compared to naïve infected or mock mice, indicating a vaccine-induced immune response at day 3 p.i. This effect was specific to the vaccine dose, as we observed no significant increase in the parenchymal CD45^+^ cell counts in 5 μg S-2P-vaccinated infected mice (**Supplemental Figure 3A**). Furthermore, the proportions of CD45^+^ parenchymal cell types varied across infection and vaccination status of mice (**Figure 2D, Supplemental Figure 3B**). The level of inflammation and proportions of immune cells in the lung parenchyma from uninfected S-2P vaccinated mice did not differ from uninfected unvaccinated mice, demonstrating that vaccination alone did not induce immune cell infiltration into the lung in this model, but rather that infection with SARS-CoV-2 was the trigger for the inflammatory response observed (**Supplementary Figure 4**).

### Monocytes and macrophages from low-dose S-2P-vaccinated mice show reduced activation and pro-inflammatory markers after B.1.351 infection compared to naïve infected mice

Mice vaccinated with 0.5 μg S-2P had a significantly lower proportion of Ly6C^hi^ inflammatory monocytes than naïve mice after infection among total CD45^+^ parenchymal cells and total monocytes (**Figure 3A**). Surface expression of the activation marker CD86 in Ly6C^hi^ and Ly6C^lo^ monocytes naïve infected mice was significantly higher compared to both mock and 0.5 μg S-2P-vaccinated infected mice (**Figure 3B-C**). Conversely, CD86 expression on Ly6C^hi^ and Ly6C^lo^ monocytes was significantly lower in 0.5 μg S-2P-vaccinated infected mice compared to naïve mice. Overall, these results reveal a distinct population of monocytes are recruited to the lungs of 0.5 μg S-2P-vaccinated mice, suggesting a different inflammatory environment in the lungs of vaccinated mice when compared to naive mice following breakthrough infection.

**Figure 3.**
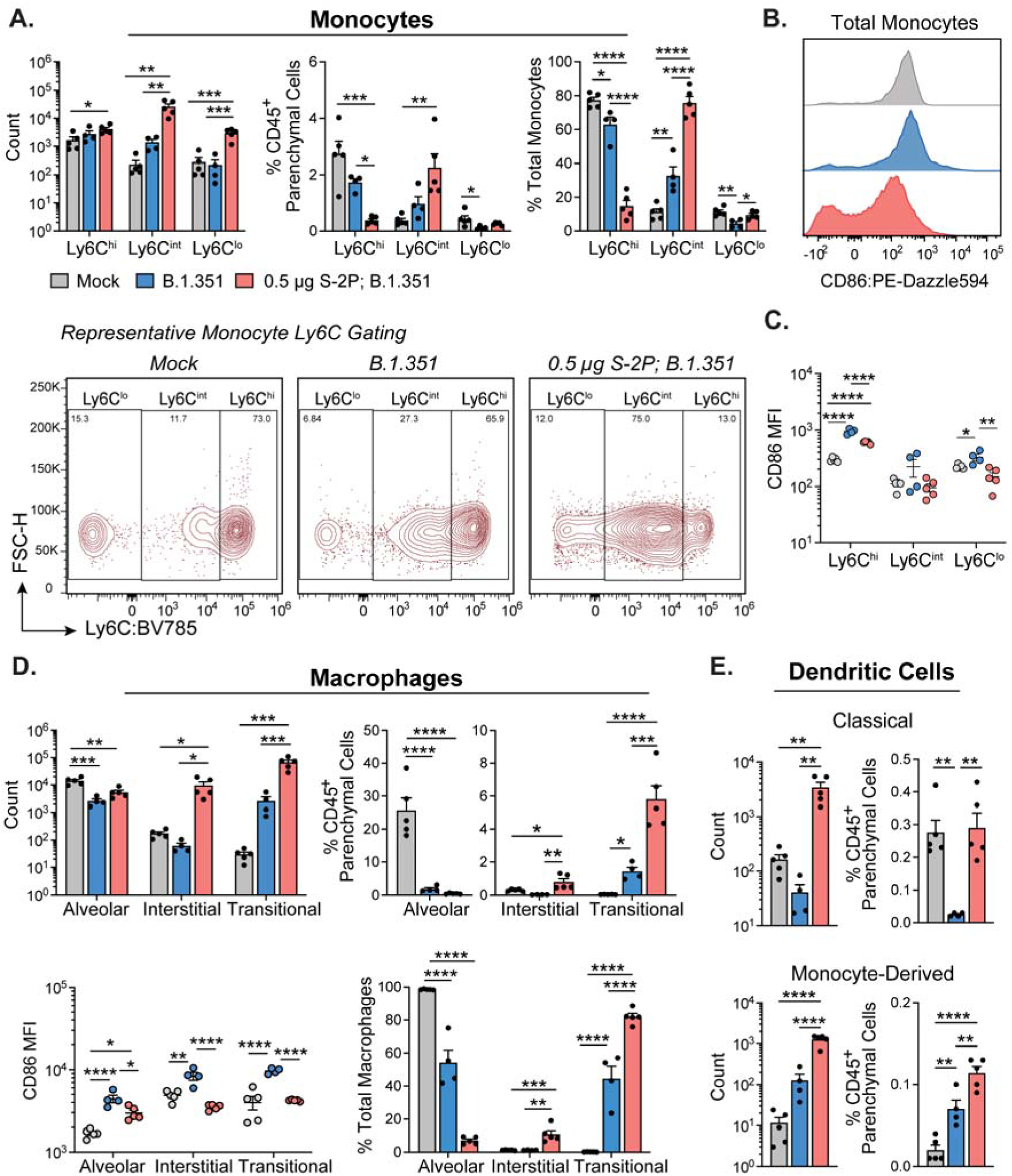
Low-dose S-2P-vaccinated B.1.351-infected mice have a unique profile of monocyte, macrophage, and dendritic cell populations in the lungs. Quantification of lung parenchymal monocytes, macrophages, and dendritic cells at day 3 p.i. (**A**) Monocyte subtypes based on Ly6C expression represented as absolute counts (top left), proportion of total CD45^+^ parenchymal cells (top middle), and proportion of total monocytes (top right). Representative gating of Ly6C^lo^ and Ly6C^hi^ monocyte subtypes (bottom). (**B**) Representative histogram of monocyte CD86 expression across groups (**C**) Mean fluorescence intensity (MFI) of CD86 expression across Ly6C monocyte populations. (**D**) Macrophage subtypes identified as alveolar, interstitial, and transitional macrophages represented as absolute counts (top left), proportion of total CD45^+^ parenchymal cells (top right), proportion of total macrophages (bottom left) and MFI of CD86 expression (bottom right). (**E**) Classical dendritic cells represented as absolute counts (top left) and proportion of total CD45^+^ parenchymal cells (top right); monocyte-derived dendritic cells represented as absolute counts (bottom left) and proportion of total CD45^+^ parenchymal cells (bottom right). Group names, sample size, and color are as follows: uninfected/mock, n=5, grey; naïve infected/B.1.351, n=4, blue; low-dose vaccinated infected/0.5 µg S-2P, B.1.351, n=5, red. Data are represented by the mean +/- the standard error of the mean. Statistical significance was determined using an unpaired one-way ANOVA with Tukey’s multiple comparisons test, and P values are represented above the bar graphs as follows: *, P < 0.05; **, P < 0.01; ***, P < 0.001; ****, P < 0.0001.

We also observed an influx of monocyte-derived macrophages (interstitial and transitional) into the lung parenchyma of 0.5 μg S-2P-vaccinated infected mice (**Figure 3D**). Proportionally, we observed a significant increase in interstitial and transitional macrophages among CD45^+^ parenchymal cells and total macrophages in the 0.5 μg S-2P-vaccinated infected mice compard to both naïve infected and mock mice. The transitional macrophages, differentiated by high expression of Ly6C on non-alveolar macrophages, were the dominant macrophage population in the lungs of 0.5 μg S-2P-vaccinated infected mice. We observed a significant reduction in alveolar macrophages in both naïve infected and 0.5 μg S-2P-vaccinated infected mice compared to mock. A reduction in tissue-resident alveolar macrophages from baseline has been previously observed following SARS-CoV-2 infection of mice and non-human primates (*19, 33, 34*). Cell surface expression of CD86 was significantly elevated in all three macrophage subtypes in the naïve infected compared to mock and 0.5 μg S-2P-vaccinated infected mice (**Figure 3D, lower left**). Only the alveolar macrophages in the 0.5 μg S-2P-vaccinated infected mice had significantly elevated CD86 expression compared to mock. The 0.5 μg S-2P-vaccinated infected mice showed similar proportion of cDCs among total CD45^+^ parenchymal cells as mock, whereas naïve infected mice exhibited a reduction in cDCs compared to mock infected and 0.5 μg S-2P-vaccinated infected mice (**Figure 3E**). There was a significant increase of moDCs in naïve infected and 0.5 μg S-2P-vaccinated infected mice compared to mock mice by both count and proportion of CD45^+^ parenchymal cells. However, there was a significantly higher proportion of moDCs in 0.5 μg S-2P-vaccinated infected compared to naïve infected mice (**Figure 3E**). We made similar observations in 5 μg vaccinated infected mice (**Supplementary Figure 3**). Taken together, these results demonstrate that vaccination status impacts the early innate immune response to B.1.351 challenge in the lungs.

### B.1.351 infections result in an influx of eosinophils in vaccinated mice

We next examined granulocyte population using the following markers to define neutrophils and eosinophils: Ly6G+ CD11b+ and CD64-CD11c-SiglecF+, respectively. In 0.5 μg S-2P-vaccinated infected mice, we found that absolute counts of both neutrophils and eosinophils were higher compared to mock and naïve infected mice (**Figure 4A, left**). However, 0.5 μg S-2P-vaccinated infected mice had a significantly higher proportion of eosinophils in the lungs among CD45^+^ parenchymal cells as compared to both mock and naïve infected mice (**Figure 4A, left; Figure 4B**). We observed no statistical difference in the proportion of eosinophils between naïve infected and mock. Conversely, naïve infected mice had a significantly higher proportion of neutrophils in the lungs compared to both mock and vaccinated infected mice. We also observed an increase in eosinophils in the lungs of 5 μg S-2P-vaccinated infected mice compared to mock and naïve infected mice, although the cell counts were lower and likely reflect low virus replication in the lungs (**Supplemental Figure 3C**). Importantly, these results were not vaccine platform-specific, as vaccination with a low-dose mRNA-1273 generating insufficient neutralizing antibody titers to control the infection resulted in an elevated influx of eosinophils after challenge with SARS-CoV-2 (**Supplemental Figure 5**). Furthermore, we observed a similar influx of eosinophils in low-dose vaccinated mice when challenged with MA-SARS-CoV-2, which is more antigenically related to the Wuhan-1 spike protein used for vaccination, though to a lesser extent (**Supplemental Figure 5)**.

**Figure 4.**
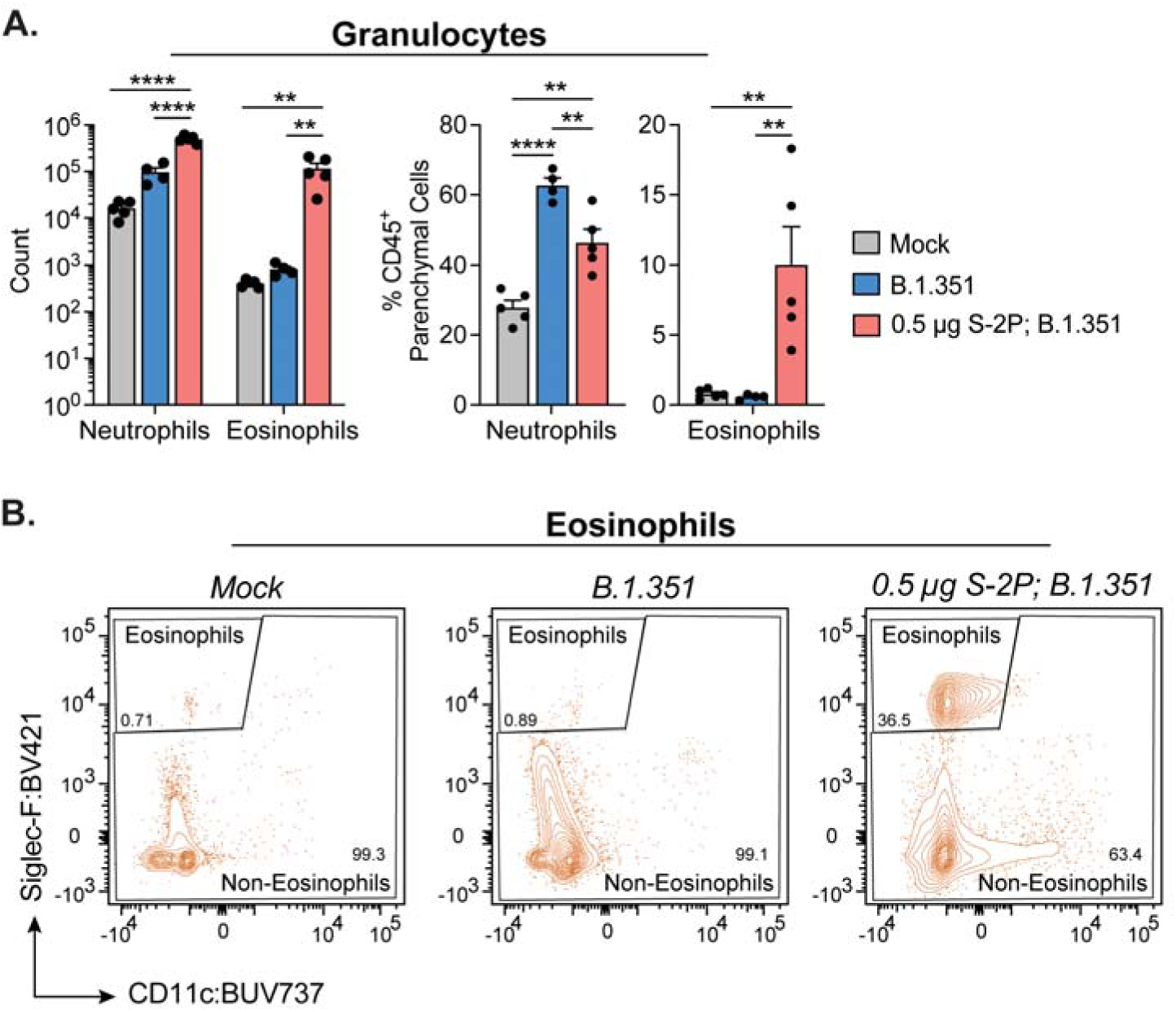
Eosinophils are a key feature of the innate immune response to B.1.351 in low-dose S-2P-vaccinated mice. Quantification of lung parenchymal granulocytes at day 3 p.i. (**A**) Granulocytes identified as neutrophils and eosinophils represented as absolute counts (left) and proportion of total CD45^+^ parenchymal cells (right). (**B**) Representative gating of eosinophils for each group. Group names, sample size, and color are as follows: uninfected/mock, n=5, grey; naïve infected/B.1.351, n=4, blue; low-dose vaccinated infected/0.5 µg S-2P, B.1.351, n=5, red. Data are represented by the mean +/- the standard error of the mean. Statistical significance was determined using an unpaired one-way ANOVA with Tukey’s multiple comparisons test, and P values are represented above the bar graphs as follows: **, P < 0.01; ****, P < 0.0001.

### SARS-CoV-2 viral RNA associates with eosinophils, monocytes, macrophages, and dead cells in low-dose S-2P-vaccinated mice at day 3 p.i

We next performed single cell RNA sequencing (scRNA-seq) to evaluate gene expression in distinct subsets of immune cells after SARS-CoV-2 infection. Parenchymal (CD45 IV-negative) cells isolated from the lungs of representative mice from the mock, naïve infected, and 0.5 μg S-2P-vaccinated infected groups (n=2 mice per group) were subjected to 10X Chromium partitioning and barcoding, library preparation, and RNA sequencing. Immune cell subsets were identified based on clustering and transcriptional profiles previously generated through scRNA-seq combined with cellular indexing of transcriptomes and epitopes (CITE-seq) (*35*). Uniform Manifold Approximation and Projection (UMAP) plots split by group (mock, naïve infected, and 0.5 μg S-2P-vaccinated infected) showing 17 subsets of CD45^+^ cells identified within the lung are shown in **Figure 5A**. Notably, there were more dead cells detected in the naïve infected mice compared to mock or 0.5 μg S-2P-vaccinated infected mice. In the 0.5 μg S-2P-vaccinated mice, we observed an increase in the eosinophil population compared to both mock and naïve infected mice, confirming our observations by flow cytometry.

**Figure 5.**
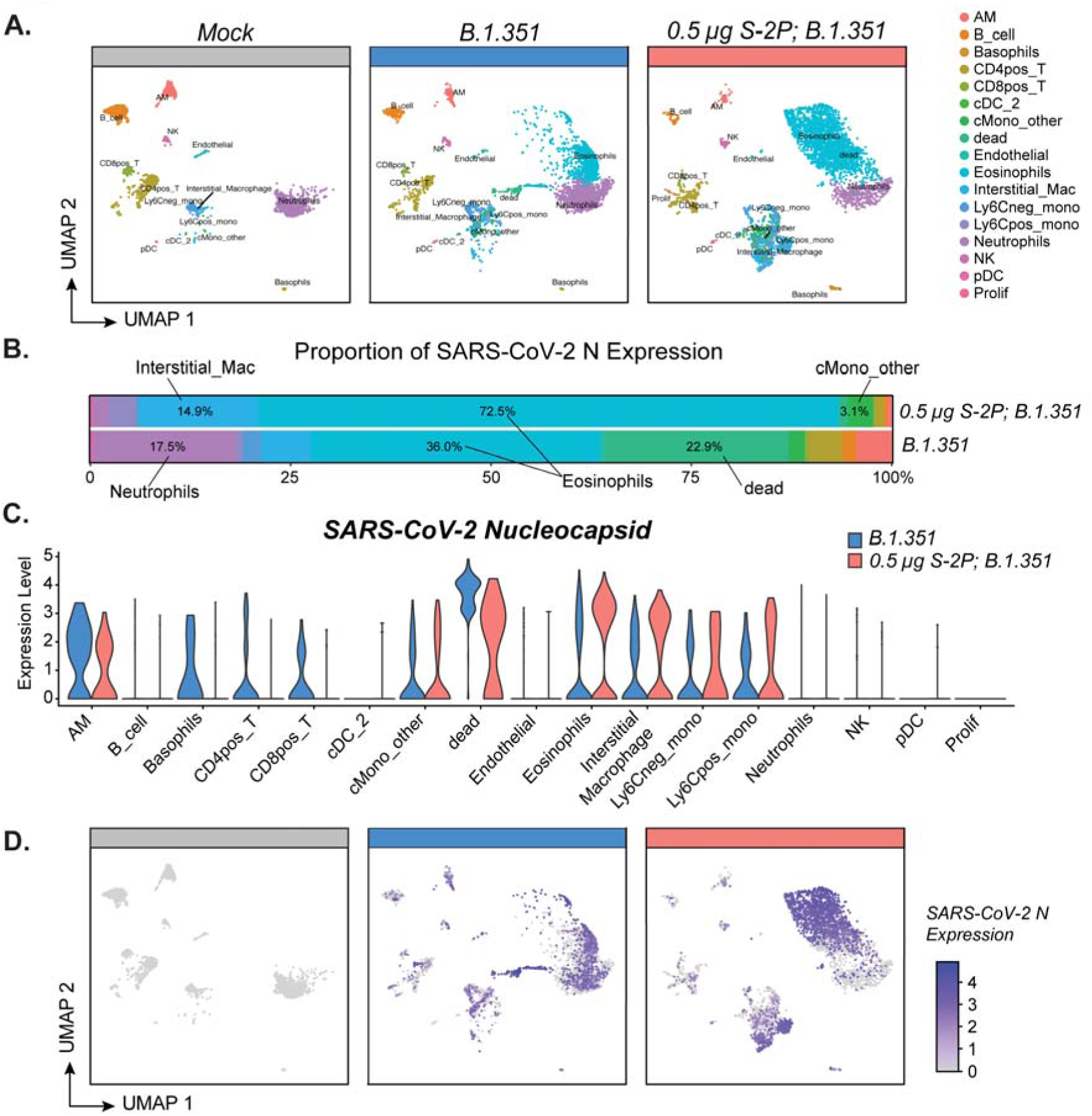
SARS-CoV-2 viral RNA is associated with eosinophils, macrophages, and monocytes in the lungs after B.1.351 infection. Single cell RNA-sequencing (scRNA-seq) analysis of parenchymal lung cells at day 3 p.i. (n=2 mice per group). (**A**) Uniform manifold approximation and projection (UMAP) plot split by group indicating the different subsets of cells identified. (**B-C**) Expression of SARS-CoV-2 N across each cell population separated by group. (**D**) UMAP featuring SARS-CoV-2 nucleocapsid (N) RNA reads within each cell. Relative expression is indicated according to the scale bar. Group names, sample size, and color are as follows: uninfected/mock, n=2, grey; naïve infected/B.1.351, n=2, blue; low-dose vaccinated infected/0.5 µg S-2P, B.1.351, n=2, red.

The SARS-CoV-2 viral RNA that was detected most commonly mapped to the nucleoprotein (N) gene or to the large open reading frame (ORF)1ab gene (**Supplemental Figure 6**). In both 0.5 μg S-2P-vaccinated and naïve infected mice, the majority of the SARS-CoV-2 N viral RNA expression in the lung parenchyma was associated with eosinophils (**Figure 5B-D**). However, the other top cell types associated with SARS-CoV-2 N viral RNA differed depending on vaccination status. Other than eosinophils, the majority of the SARS-CoV-2 N viral RNA was most highly associated with interstitial macrophages and monocytes in 0.5 μg S-2P-vaccinated mice, whereas in the naïve mice SARS-CoV-2 N viral RNA was most highly associated with dead cells and neutrophils (**Figure 5B**). SARS-CoV-2 N viral RNA was also found in the cluster of dead cells that was composed of cells predominantly from naïve infected mice (**Figure 5C**).

### Monocyte and interstitial macrophage signaling contributes to the immune response observed in low-dose S-2P-vaccinated mice after infection

We next analyzed the transcriptional signature of monocytes and interstitial macrophages (**Figure 6**). Monocytes and interstitial macrophages from naïve infected and 0.5 μg S-2P-vaccinated infected mice were transcriptionally distinct, localizing in different spatial locations on the UMAP and expressing different gene markers (**Figure 6B-C**). Particularly, genes associated with M2-like macrophages (*Arg1*, *Chil3*) were found to be defining markers for the interstitial macrophages and monocytes in 0.5 μg S-2P-vaccinated mice (**Figure 6D**). While a higher expression level of SARS-CoV-2 N was observed in interstitial macrophages and monocytes from 0.5 μg S-2P-vaccinated mice compared to naïve, there was a reduced enrichment in the interferon (IFN) alpha response pathway as determined by gene set enrichment analysis (GSEA) (**Figure 6E-F**) (*36*). We also observed high expression of *Ccl24* in the 0.5 μg S-2P-vaccinated mice, the gene for eotaxin-2, which is a chemokine responsible for recruiting eosinophils to the site of infection (**Figure 6G**) (*37*). Furthermore, in the 0.5 μg S-2P-vaccinated infected mice, monocytes and interstitial macrophages were enriched for granulocyte chemotaxis pathway by GSEA compared to naïve infected (**Figure 6H**). This suggests that the interstitial macrophages and monocytes are involved in recruiting eosinophils to the lung parenchyma after breakthrough infection via *Ccl24* expression.

**Figure 6.**
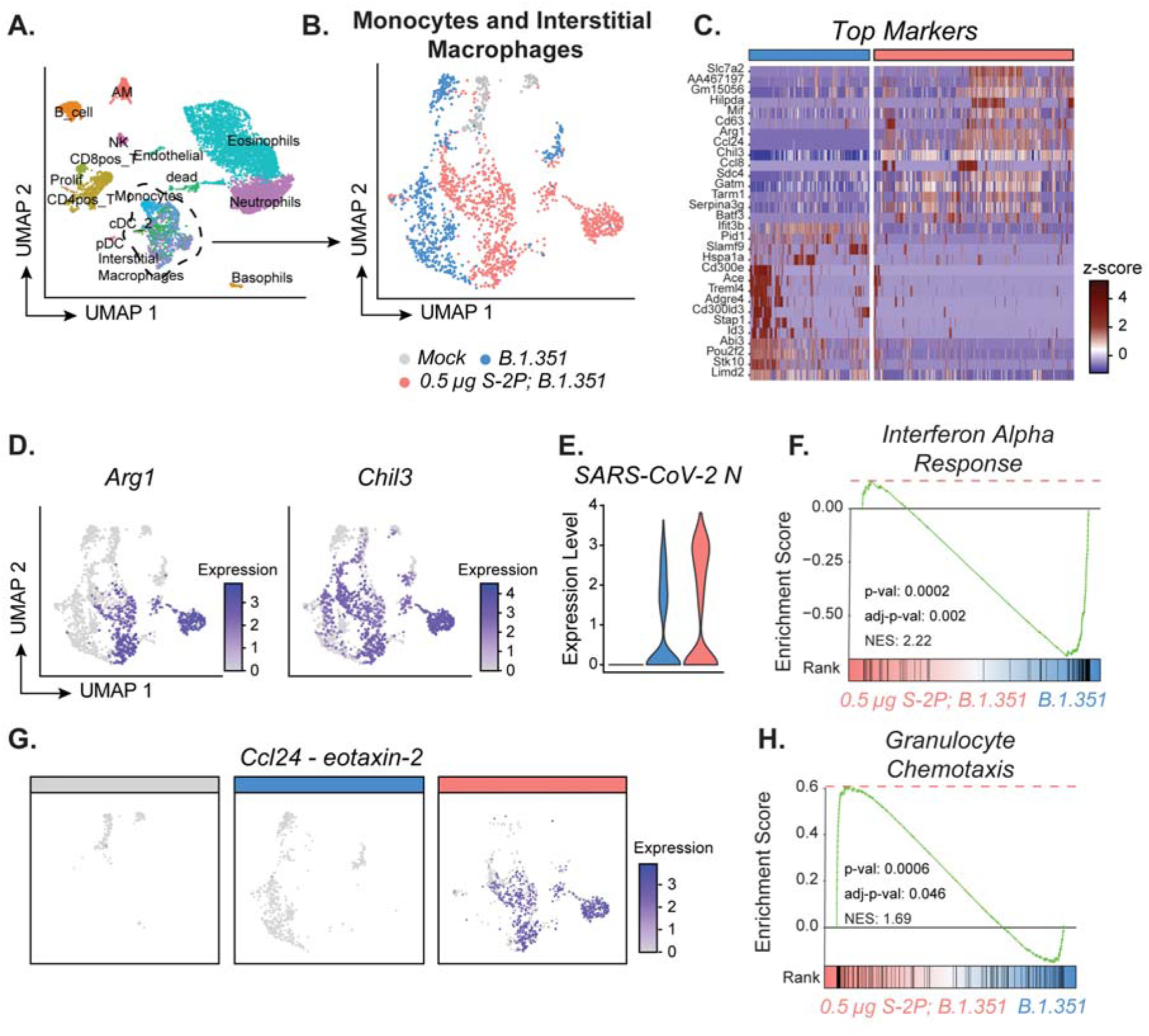
Lung monocytes from low-dose S-2P-vaccinated B.1.351 infected mice express features for granulocyte chemotaxis. Single cell RNA-sequencing (scRNA-seq) analysis of parenchymal lung cells at day 3 p.i. (n=2 mice per group). (**A**) UMAP of all samples combined displaying the different cell types identified. (**B**) UMAP of the interstitial macrophages and monocytes subsetted from (**A**) and color coded based on group: (uninfected) mock = grey, (naïve infected) B.1.351 = blue, (vaccinated infected) 0.5 µg S-2P, B.1.351 = red. (**C**) Heatmap featuring the top markers for the interstitial macrophages/monocytes between B.1.351 and 0.5 µg S-2P, B.1.351 determined by differential gene expression analysis. Rows are genes and columns are individual cells organized by group. Data are represented as row z-scores. (**D**) UMAP featuring *Arg1* and *Chil3* expression in interstitial macrophages and monocytes. (**E**) SARS-CoV-2 N expression in the subset of interstitial macrophages and monocytes across mock, B.1.351, and 0.5 µg S-2P, B.1.351. (**F**) Gene set enrichment analysis (GSEA) plot for Hallmark Interferon Alpha Response pathway between B.1.351 = blue, 0.5 µg S-2P, B.1.351 = red. (**G**) UMAP split based on group featuring Ccl24 expression in interstitial macrophages and monocytes. (**H**) GSEA plot for GOBP Granulocyte Chemotaxis pathway (GO:0071621) between B.1.351 = blue, 0.5 µg S-2P, B.1.351 = red. Group names, sample size, and color are as follows: uninfected/mock, n=2, grey; naïve infected/B.1.351, n=2, blue; low-dose vaccinated infected/0.5 µg S-2P, B.1.351, n=2, red.

### Eosinophils from low-dose S-2P-vaccinated infected mice are phenotypically distinct from eosinophils from mock and naïve infected mice

We next explored the transcriptional signatures of granulocytes by scRNA-seq (**Figure 7A**). Granulocytes were subsetted from the other cell types (**Figure 7B**). Similar to the results by flow cytometry, we observed a greater proportion of neutrophils in the naïve infected mice compared to the 0.5 μg S-2P-vaccinated infected mice, whereas a greater proportion of eosinophils was observed in the 0.5 μg S-2P-vaccinated infected mice (**Figure 7C**). Granulocytes divided into 5 transcriptionally distinct clusters (**Figure 7D-E**). We identified 3 major clusters of eosinophils labeled 0, 2, and 4 and segregated based on naïve or 0.5 μg S-2P-vaccinated infected mice. Eosinophils from naïve infected mice were found predominantly in clusters 2 and 4. Cluster 0 was almost exclusively occupied by eosinophils from 0.5 μg S-2P-vaccinated mice (**Figure 7E**). Interestingly, the top markers for cluster 4 were SARS-CoV-2 viral genes and included ORF1ab, ORF8, Spike (S), Membrane (M), ORF7a, ORF3a, Envelope (E), and ORF6 (**Figure 7D**). Despite a high level of viral RNA in the cluster 0, the defining markers were not SARS-CoV-2 viral genes. A notable top marker for cluster 0, which was composed of 0.5 μg S-2P-vaccinated eosinophils, was *Cstb* (**Figure 7D, G**). *Cstb* is the gene for Cystatin B, an inhibitor of Cysteine protease B, which has been shown to be involved in SARS-CoV-2 entry via the endosome (*38*). Despite the higher level of SARS-CoV-2 viral RNA in the 0.5 μg S-2P-vaccinated eosinophils, there was no difference in the response to IFN-alpha between 0.5 μg S-2P-vaccinated or naïve infected mice by GSEA (**Supplemental Figure 7**). There was limited overlap in expression between the top transcriptional markers for eosinophils from naïve infected mice compared to the 0.5 μg S-2P-vaccinated infected (**Figure 7H**). Eosinophils can be influenced by different cytokine milieus, including IFN-γ and IL-4 (*39–41*). To better understand what defines these transcriptional differences, we generated modules from previously published bulk RNA-seq data from eosinophils isolated from mice and measured enrichment in these modules across our eosinophils (**Figure 7I**) (*41*). The Asthma module was derived from top DEGs from lung eosinophils isolated from a mouse model of asthma. The IFN-γ and IL-4 modules were derived from top DEGs from eosinophils isolated from peritoneal cavity and treated with IFN-γ or IL-4, respectively. Eosinophils from 0.5 μg S-2P-vaccinated infected mice showed enrichment in the IFN-γ module, but not the Asthma or IL-4 modules. By flow cytometry, eosinophils from naïve infected mice showed upregulated CD86 and MHC-I, whereas those from 0.5 μg S-2P-vaccinated infected mice expressed similar levels as mock eosinophils (**Figure 7J**). We observed the same expression level of MHC-II, which has been shown to be upregulated in response to allergen challenge, and CD107a, a marker for degranulation, among mock, naive infected, and vaccinated infected mice (**Supplemental Figure 8**). The SSC-A, a marker of granularity, of eosinophils from 0.5 μg S-2P-vaccinated infected mice was elevated compared to mock and naïve infected mice, which has been linked to viability and protease activity (**Figure 7J**) (*42*). Taken together, these data demonstrate that eosinophils associated with SARS-CoV-2 breakthrough infections have a unique phenotype, including a transcriptional signature for an IFN-γ response, low activation marker expression, and high SSC-A in our model.

**Figure 7.**
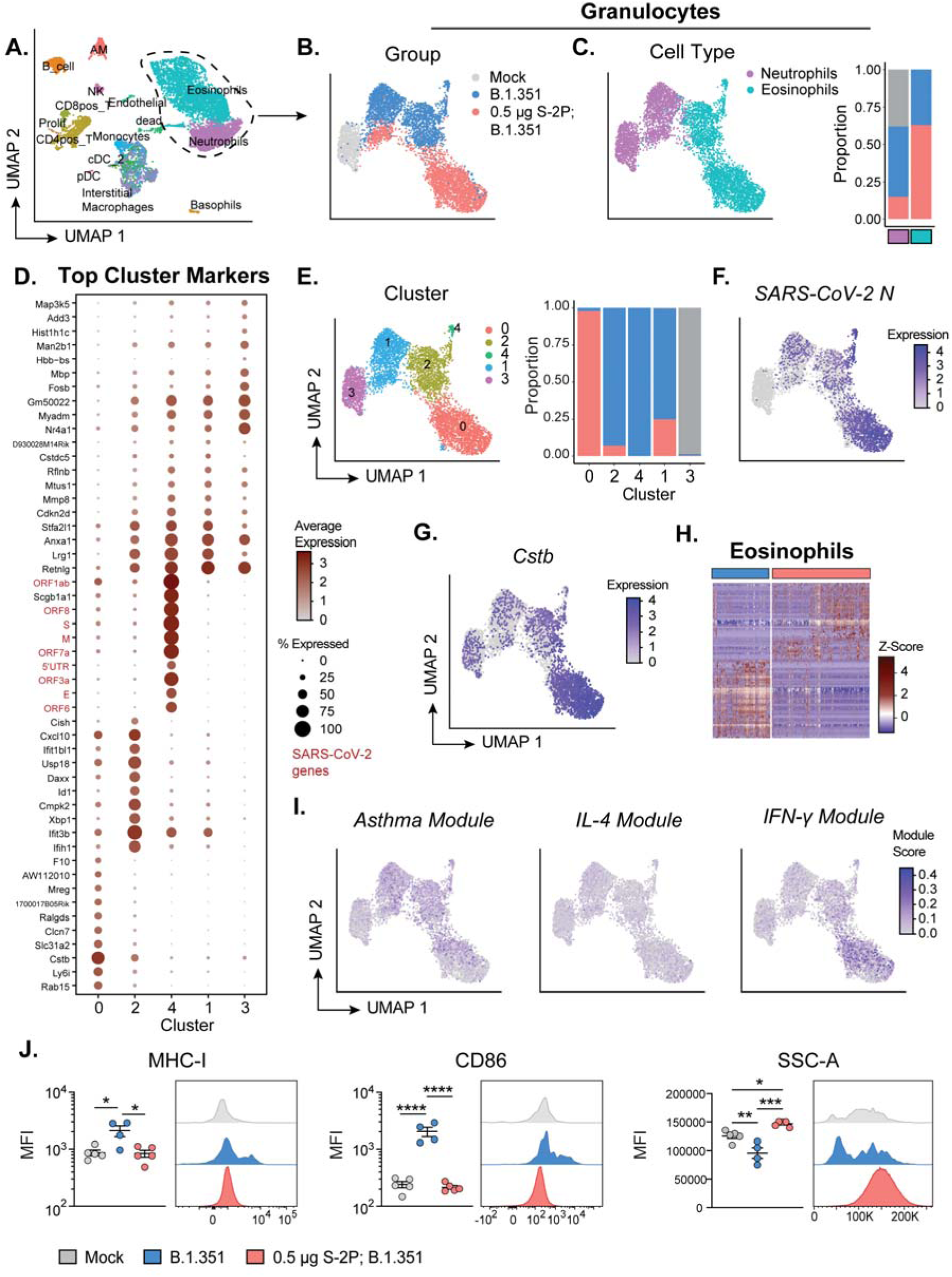
Eosinophils from low-dose S-2P-vaccinated infected mice are phenotypically distinct from eosinophils from mock and naïve infected mice. Single cell RNA-sequencing (scRNA-seq) analysis of parenchymal lung cells at day 3 p.i. (n=2 mice per group). (**A**) UMAP of all samples combined displaying the different cell types identified. (**B**) UMAP of the granulocytes identified as neutrophils and eosinophils subsetted from (**A**) displaying the group of each cell: (uninfected) mock = grey, (naïve infected) B.1.351 = blue, (vaccinated infected), B.1.351 = red. (**C**) UMAP (left) of the subsetted granulocytes identified as neutrophils and eosinophils displaying the specifc cell type: neutrophil = purple and eosinophil = teal. Frequency distribution for the groups that make up the neutrophil and eosinophil populations (right). (**D**) Top markers for each of the 5 clusters identified among the granulocytes. (**E**). UMAP (left) of the subsetted granulocytes displaying the assigned cluster. Frequency distribution for the groups that make up each granulocyte cluster (right). (**F-G**) UMAPs of the subsetted granulocytes featuring *SARS-COV-2 N* (**F**) and *Cstb* (**G**) expression. (**H**) Heatmap featuring the top markers for the eosinophils between B.1.351 and 0.5 µg S-2P, B.1.351 determined by differential gene expression analysis. Rows are genes and columns are individual cells organized by group. Data are represented as row z-scores. (**I**) UMAPs of the subsetted granulocytes featuring scores for transcriptional modules for asthma, IL-4 treatment, and IFN-gamma treatment. (**J**) Flow cytometry analysis of lung parenchymal eosinophil cell surface expression at 3□days p.i. Plots displaying mean fluorescence intensity (MFI) (left). Representative histograms of MHC-I, and CD86 expression and the SSC-A profile (right). Group names, sample size, and color are as follows: uninfected/mock, n=2 (scRNA-seq) or n=5 (flow cytometry), grey; naïve infected/B.1.351, n=2 (scRNA-seq) or n=4 (flow cytometry), blue; low-dose vaccinated infected/0.5 µg S-2P, B.1.351, n=2 (scRNA-seq) or n=5 (flow cytometry), red. Data are represented by the mean +/- the standard error of the mean. Statistical significance was determined using an unpaired one-way ANOVA with Tukey’s multiple comparisons test, and P values are represented above the bar graphs as follows: *, P < 0.05; **, P < 0.01; ***, P < 0.001; ****, P < 0.0001.

### Eosinophil depletion reduces lung protection from B.1.351 challenge of low-dose S-2P-vaccinated mice

To determine a role of eosinophils during SARS-CoV-2 breakthrough infection, we depleted eosinophils in 0.5 μg S-2P-vaccinated mice with an α-CCR3 (receptor for eotaxin-2) antibody on days -1, 0, 1, and 2 p.i. (**Figure 8A**). 0.5 μg S-2P-vaccinated mice were injected with an antibody of the same isotype as the α-CCR3 antibody on the same schedule as a control. Mice were challenged with B.1.351 and their lungs were harvested at 3 days p.i. for analysis. We observed that the α-CCR3 mice could be stratified based on two different levels of eosinophil depletion: less than or greater than 5-fold depletions (**Figure 8B**). Pre-challenge FRNT_50_ titers against B.1.351 were no different across the treatment groups, and thus eosinophil depletion efficiency was not impacted by neutralizing antibody levels (**Figure 8C**). The mice with >5x eosinophil depletion in the lung lost weight similar to naïve infected mice (**Figure 8D**). However, mice with <5x eosinophil depletion showed weight loss similar to isotype control infected mice. We proceeded with analysis of mice with >5x eosinophil depletions in the α-CCR3 mice. We observed a significant increase in infectious virus as determined by plaque assay in the α-CCR3 mice compared to the isotype control (**Figure 8E**). Furthermore, the distribution of SARS-CoV-2 as determined by N antigen staining in the lungs changed upon eosinophil depletion (**Figure 8F**). Viral antigen staining was restricted to the bronchial lumen in 0.5 μg S-2P-vaccinated mice treated with the isotype control, whereas viral antigen was found in the bronchial lumen, lining the bronchial epithelium, and spread throughout the alveoli in 0.5 μg S-2P-vaccinated mice treated with α-CCR3. Genomic viral RNA (RdRp) as measured by qPCR in lung lysates negatively correlated with eosinophil counts across all infected mice (**Figure 8G**).

**Figure 8.**
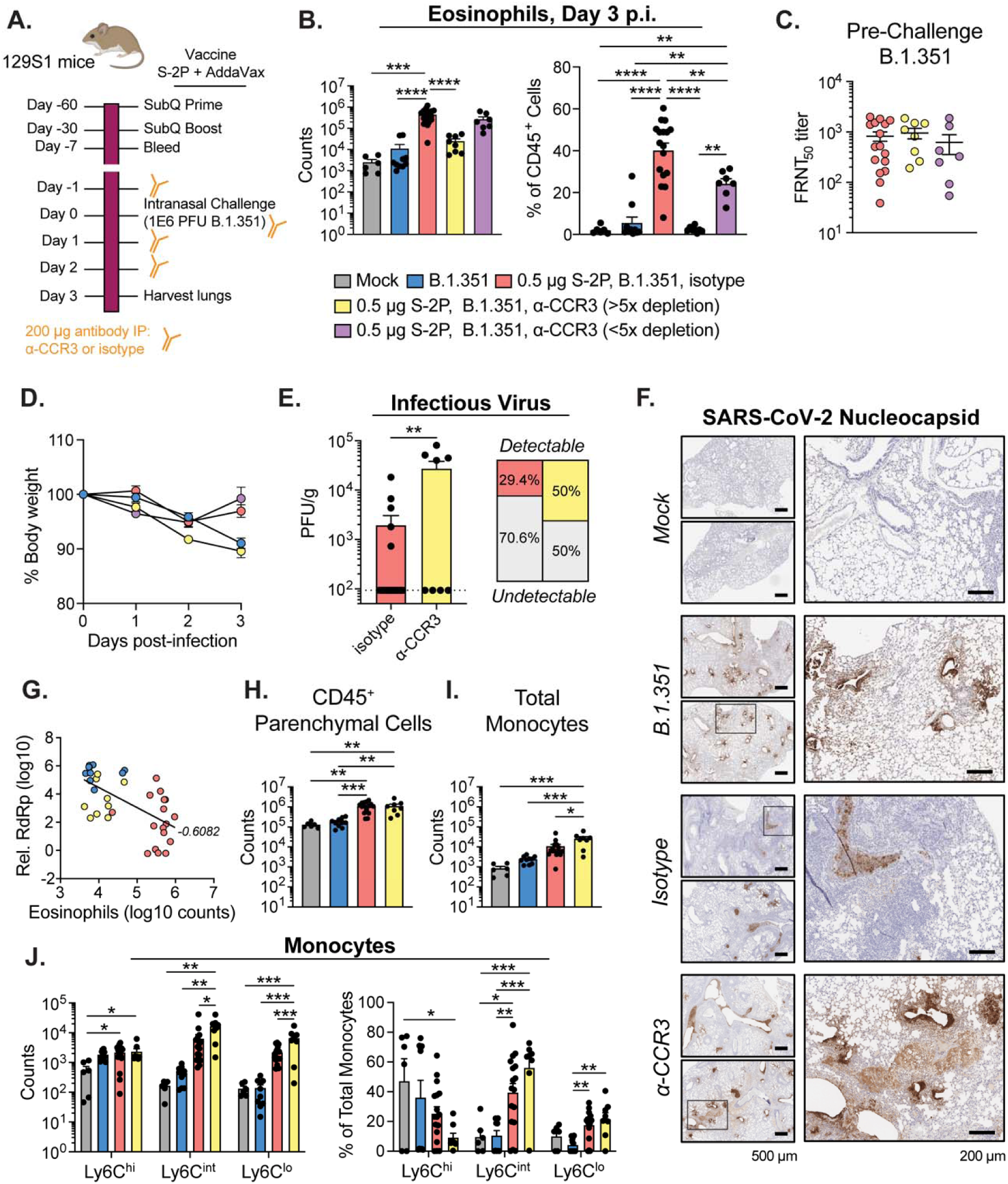
Eosinophils provide protection during B.1.351 infection in vaccinated mice. (**A**) Experimental schematic detailing vaccination of 129S1/SvImJ mice with S-2P, eosinophil depletion schedule with anti-CCR3, and subsequent challenge with B.1.351. (**B**) Quantification of lung parenchymal eosinophils represented as absolute counts (left) and proportion of CD45^+^ parenchymal cells of at day 3 p.i. (**C**) Live virus serum FRNT_50_ titers against B.1.351 prior to challenge in mice vaccinated with 0.5 µg S-2P. (**D**) Body weight loss as a percentage of day of challenge for mice infected with B.1.351. (**E**) B.1.351 viral loads in the lungs as measured by plaque assay at day 3 p.i. Percentages represent the frequency of samples with detectable infectious virus by plaque assay. (**F**) Representative images of histopathological staining for SARS-CoV-2 nucleoprotein (N) of mouse lungs harvested at day 3 p.i. Scale bars are 500 µm (left) and 200 µm (right). (**G**) Correlation analysis between relative expression of genomic viral RNA (RdRp) determined by RT-qPCR and eosinophil counts determined by flow cytometry. Line represents simple linear regression and statistic is the Spearman correlation coefficient, r. (**H**) Absolute counts of CD45^+^ lung parenchymal cells and (**I**) total monocytes in the lung parenchyma. (**J**) Quantification of lung parenchymal monocytes identified as Ly6C^lo^, Ly6C^int^, and Ly6C^hi^ represented as absolute counts (left) and proportion of total monocytes (right). Group names, sample size, and color are as follows: uninfected/mock, n=6, grey; naïve infected/B.1.351, n=10, blue; 0.5 µg S-2P, B.1.351, isotype, n=17, red; 0.5 µg S-2P, B.1.351, anti-CCR3 (>10X eosinophil depletion), n=8, yellow; 0.5 µg S-2P, B.1.351, anti-CCR3 (<10X eosinophil depletion), n=7, purple. Data are represented by the mean +/- the standard error of the mean. Statistical significance was determined using an unpaired Student’s *t* test or one-way ANOVA with Tukey’s multiple comparisons test, and P values are represented above the bar graphs as follows: *, P < 0.05; **, P < 0.01; ***, P < 0.001; ****, P < 0.0001. Created with BioRender.com.

Despite a 1-log fold reduction in eosinophil counts there was no change in the total number of CD45^+^ parenchymal cells in the α-CCR3 mice compared to the isotype control, indicating that there was compensation for the lack of eosinophils by recruitment of other cell types (**Figure 8H**). Notably, a significant increase in total monocytes was observed in 0.5 μg S-2P-vaccinated mice treated with α-CCR3 compared mock, naïve infected, and 0.5 μg S-2P-vaccinated mice treated with isotype (**Figure 8I**). Furthermore, there was an increase in both Ly6C^int^ and Ly6C^lo^ monocytes in the α-CCR3 mice compared to the isotype control (**Figure 8J**). These results indicate that the presence of eosinophils impact the broader inflammatory mileu during breakthrough infections.

## Discussion

To understand the immune response during SARS-CoV-2 vaccine breakthrough infection, we modeled waning immunity by vaccinating mice with a low-dose of the ancestral S-2P protein subunit vaccine allowing us to establish a vaccine-mismatch SARS-CoV-2 infection in the lungs with B.1.351. Breakthrough infections resulted in lower viral loads in the lungs and reduced weight loss compared to naïve mice with no prior SARS-CoV-2 antigen exposure. Using this model, we found that the innate immune response in the lungs during the acute phase of the breakthrough infection was distinct from the response to infection in a naïve mouse. We observed an increase in monocytes, macrophages, dendritic cells, eosinophils, and neutrophils in the breakthrough infection model compared to mock or naïve infected mice. We found that the innate immune response to SARS-CoV-2 breakthrough infection was overall less inflammatory compared to naïve infections based on the proportions and phenotypes of the immune cells present in the lung parenchyma. Breakthrough infections were characterized by a high proportion of eosinophils, monocytes and macrophages with reduced activation marker expression compared to naïve infections. In contrast, naïve infections resulted in a high proportion of neutrophils and activated monocytes and macrophages. We found that eosinophils were protective in the context of SARS-CoV-2 breakthrough infections, as depletion of eosinophils in low-dose vaccinated mice had greater weight loss, lung viral load, and viral dissemination throughout the lungs compared to isotype control after challenge.

We observed that SARS-CoV-2 breakthrough infections were defined by an influx of eosinophils into the lung parenchyma early after infection. The breakthrough infection-associated eosinophils were phenotypically distinct from eosinophils found in the lungs during naïve or mock infections. Eosinophils are commonly known for their roles in Th2-skewed immune responses that mediate host defense against parasitic infections and the pathology of allergic asthma (*43*). There is growing evidence that eosinophils can also be important players in the host antiviral immunity to respiratory viral infections, such as respiratory syncytial virus (RSV), parainfluenza virus, and influenza virus (*44–48*). The antiviral action of eosinophils has been observed both indirectly by antigen presentation and activation of T cells and directly by cationic proteins, RNases, and reactive oxygen and nitrogen species (*44–50*). However, there remains substantial debate as to whether eosinophils contribute to pulmonary pathology during respiratory viral infections, promote viral clearance, or both (*44, 51–53*). In hamsters, eosinophils were suggested to be present in the lungs during SARS-CoV-2 vaccine breakthrough infections with WA-1 and the Alpha variant (*54*). Recently, Chang *et al.* reported non-pathological lung eosinophilia in response to an influenza vaccine breakthrough infection in mice, whereby there was no increased morbidity in the form of weight loss and minimal inflammatory cytokine and chemokine expression in the lungs (*55*). We also reported minimal weight loss during SARS-CoV-2 breakthrough infection and the associated eosinophils did not bear transcriptional similarity with eosinophils from an asthma model, suggesting that the eosinophilia that we observed might also be non-pathological. It is possible that eosinophils contribute to either pathology or protection depending on a number of factors such as the specific virus and the corresponding eosinophil phenotype, which in turn may be influenced by the inflammatory environment in the lungs.

Eosinophils have been a hallmark of pathogenicity in the context of breakthrough infections and a warning sign for potentially harmful vaccine formulations in pre-clinical studies (*56–58*). Most notably, in the 1960s, vaccination with formalin-inactivated RSV in children led to enhanced disease including hospitalization, pneumonia, and mortality in two children when they acquired a natural RSV infection (*59–61*). Eosinophils were noted in the lungs of one child who perished (*61*). Vaccine-associated enhanced disease has since been studied in small animal models and is defined by a Th2-skewed inflammatory response including the presence of eosinophils in the lungs (*58*). Depletion of eosinophils in mouse models of RSV vaccine-enhanced disease have been shown to result in higher RSV viral titers indicating some level of protection by eosinophils, yet different measures have been used to assess their role in enhanced disease leading to conflicting conclusions (*52, 53, 62*). Castilow *et al.* reported weight loss, clinical illness, and airway resistance as determined by whole body plethysmography after RSV infection even in the absence of eosinophils, suggesting that eosinophils are not the drivers of vaccine-enhanced disease (*58, 62*). Our data supports the hypothesis that eosinophils can play a protective and antiviral role during a respiratory vaccine breakthrough infection and highlights the importance of investigating the quality of the eosinophilic response along with the quantity. Further work is needed to determine the role that eosinophils play during vaccine breakthrough infections, their direct impact on lung function, and if there are specific phenotypes that define pathogenic or protective eosinophils for different respiratory viruses.

Depletion of eosinophils in vaccinated mice correlated with increased viral burden in the lungs, viral spread into the alveolar space, and increased morbidity as determined by weight loss. We observed a high level of viral RNA associated with the breakthrough infection-associated eosinophils by scRNA-seq which could contribute to their antiviral state. It is unclear if the eosinophils were infected with SARS-CoV-2 or had internalized virus particles or infected cells by phagocytosis. A defining marker of breakthrough infection-associated eosinophils was their expression of *Cstb*, which is an inhibitor of a known SARS-CoV-2 entry factor. These data suggest that the protective role that eosinophils play during SARS-CoV-2 vaccine breakthrough in the lung may be through direct antiviral action. Oyesola *et al.* reported that recent exposure to lung migrating helminth infection protected against SARS-CoV-2 challenge in mice, where they noted that eosinophils were present in the lungs at higher levels at the time of infection (*63*).

They found that alternatively activated (M2-like) interstitial macrophages mediated viral clearance via activation of CD8+ T cells on day 7 p.i. In our SARS-CoV-2 breakthrough infection model, we also observed an M2-like phenotype among interstitial macrophages and monocytes including high expression of *Arg1*, *Chil3*, and *Ccl24*, similar to the phenotype from the helminth model. We hypothesize that the monocytes and interstitial macrophages recruit the eosinophils to the lung during breakthrough infection based on their expression of *Ccl24.* We observed an increased number of monocytes in the lung parenchyma of vaccinated mice with depleted eosinophils after challenge compared to the isotype control, indicating that eosinophils also influence the broader inflammatory milieu which may contribute to protection. Future studies should be aimed at investigating the antiviral activity of eosinophils and the interplay between interstitial macrophages, monocytes, and eosinophils during SARS-CoV-2 breakthrough infections.

Eosinopenia (absent or reduced circulating eosinophil counts) has been demonstrated as an independent marker of severe COVID-19 and mortality in humans (*64–67*). Further, recovered eosinophil counts in hospitalized patients that were admitted with eosinopenia has been associated with improved outcomes for COVID-19. Interestingly, individuals with asthma have been reported to have reduced severity and mortality from COVID-19 (*68–70*). Although there are several possible reasons for this effect, one hypothesis has been that there is antiviral activity being exerted by the eosinophils which are present in the lung at higher baseline levels in patients with asthma (*71, 72*). In this study, the eosinophils in the lung parenchyma observed during SARS-CoV-2 breakthrough infection did not bear transcriptional similarity to the eosinophils from a previously published mouse model of asthma (*41*). However, it is important to note that the asthma model reference was not challenged with a respiratory viral infection, which would likely change the transcriptional state of the asthma-associated eosinophils. Further studies are warranted to understand the contribution of eosinophils in protection against SARS-CoV-2 in asthmatic and healthy individuals.

In summary, we report a mouse model to study vaccine breakthrough infection. We used this model to show that breakthrough infection leads to the recruitment of eosinophils which contribute to protection against SARS-CoV-2 replication to the lungs. These results highlight the critical role for the innate immune response in vaccine mediated protection against SARS-CoV-2.

## Acknowledgements

This work was supported in part by grants (P51 OD011132 to Emory University) from the National Institutes of Health (NIH), Emory Executive Vice President for Health Affairs Synergy Fund award, the I3 Synergy Awards, COVID-19 cures, CEIRR HHSN272201400004C, CEIRR (75N93021C00017), and the Systems Immunology Research at Emory University (P30AI050409). This work was also supported in part by the Division of Intramural Research of NIAID/NIH. The Flow Cytometry Core is generously supported by Emory CFAR, Emory Vaccine Center and Emory National Primate Research Center of Emory University.

## Author contributions

K.M.M., S.L.F, M.S.S. contributed to the acquisition, analysis, and interpretation of the data, as well as the conception and design of the work, and writing of the manuscript. M.K. contributed to the acquisition and analysis of the data. E.J.E., K.A.F., M.K., J.W.V., M.E., A.M., B.W., S.L., A.M., R.A.S., R.R.A., S.E.B. contributed to the acquisition of the data. M.E.W. contributed to the analysis and interpretation of the data. J.E.K., V.D.M, and A.G. contributed to the interpretation of the data and conception and design of the work.

## Methods

### Viruses and cells

Vero E6 cells stably expressing human transmembrane serine protease 2 (TMPRSS2), hereafter called Vero E6-TMPRSS2 cells, were generated and cultured as previously described (Edara, *et al*., *NEJM* 2021). Vero E6 cells stably expressing TMPRSS2 and human angiotensin-converting enzyme 2 (hACE2), hereafter called Vero E6-TMPRSS2-hACE2 cells, were kindly provided by Dr. Barney Graham (Vaccine Research Center, NIH, Bethesda, Maryland, USA) and were generated and cultured as previously described (Chen *et al*., *Nat Med* 2021). The MA-SARS-CoV-2 virus was generated by engineering four coding mutations (NSP6 L37F, NSP10 P87S, S N501Y, and N D128Y) into an infectious clone (ic)SARS-CoV-2 backbone based on the original WA-1 isolate. The resulting virus was passaged 20 times in Balb/c mice, followed by deep sequencing, which identified three additional acquired mutations (S T417N, S H655Y, and E E8V). This virus was passaged once in Vero E6 cells to generate a working stock of MA-SARS-CoV-2 virus (Muruato, *et al., PLoS Biology* 2021). The B.1.351 variant isolate was kindly provided by Dr. Andy Pekosz (Johns Hopkins University, Baltimore, Maryland, USA) and was propagated once in Vero E6-TMPRSS2 cells to generate a working stock. All viruses used in this study were deep sequenced and confirmed as previously described (Edara, *et al*., *NEJM* 2021).

### Vaccination and infection of mice with SARS-CoV-2

Male or female 129S1/SvImJ mice were purchased from Jackson Laboratories at around five to six weeks of age. At eight weeks old, mice were vaccinated with either prefusion-stabilized protein subunit vaccine based on the Wuhan-1 SARS-CoV-2 isolate (S-2P) through subcutaneous injection of either 0.5 µg or 5 µg of protein formulated with AddaVax (Invivogen, #vac-adx-10) or mRNA-1273 (WA-1 matched spike) through intramuscular injection of either 0.1 µg or 1 µg. The second dose was administered four weeks later and was the same amount as the first dose for each mouse. After at least 30 days following the second dose of vaccine, mice were transferred to an animal biosafety level 3 (ABSL-3) facility and infected with SARS-CoV-2. Briefly, stock MA-SARS-CoV-2 or B.1.351 virus was diluted in sterile saline (Medline Industries, #RDI30296). Mice were anesthetized with isoflurane and infected i.n. with 50 microliters (µL) of virus solution (1 x 10^6^ PFU) per mouse. Mice were monitored daily for weight loss. For antibody-mediated depleteion of eosinophils, on days-1, 0, 1, and 2 p.i. mice were administered 0.2 mg anti-CCR3 antibody (BioXCell, clone 6S2-19-4, #BE0316) or 0.2 mg isotype control (BioXCell, Rat IgG2b, κ clone LTF-2, #BE0090) in saline by the intraperitoneal (i.p.) route. All experiments adhered to protocols approved by the Emory University Institutional Animal Care and Use Committee and were carried out under approved biosafety conditions.

### Quantification of infectious virus

At indicated days p.i., mice were euthanized via isoflurane overdose, and lung tissue was collected in 2 mL tubes containing ceramic beads (Omni International, #19-627) filled with Hank’s Balanced Salt Solution (HBSS; Sigma-Aldrich, #55021C-1000ML) supplemented with 1% fetal bovine serum (FBS; Corning, #35-011-CV). Tissue was homogenized using an Omni Bead Ruptor 24 (Omni International; 5.15 ms, 15 seconds) and then centrifuged to remove debris. To perform plaque assays, 10-fold serial dilutions of viral supernatant in serum-free Dulbecco’s Modified Eagle Medium (DMEM; Lonza, #12-614Q) were overlaid on Vero E6-TMPRSS2-hACE2 confluent monolayers and adsorbed for one hour at 37°C and 5% carbon dioxide (CO_2_). After adsorption, 0.6% immunodiffusion-grade agarose (MP Biomedical, #952014) in DMEM (Millipore Sigma, #SLM-202-B) supplemented with 5% FBS, 1% penicillin/streptomycin (Gibco, #15140163), 1% 100 mM sodium pyruvate (Corning, #25-000-CI), 1% 100X MEM Nonessential Amino Acids (Corning, #25-025-CI), 1% 1M HEPES (Corning, #25-060-CI) and 120 mM sodium bicarbonate was overlaid, and cultures were incubated until plaques were visible (about 48 hours) at 37°C and 5% CO_2_. Agarose plugs were removed, and cells were fixed and stained with solution containing final concentrations of 2% paraformaldehyde (PFA; Electron Microscopy Sciences, #15713-S) in DPBS, 10% methanol (VWR, #BDH1135-4LG) in water, and 0.05% crystal violet (Acros Organics, #405831000).

### Quantitative reverse transcription polymerase chain reaction (RT-qPCR)

At indicated days p.i., mice were euthanized via isoflurane overdose, and lung and nasal turbinate tissues were collected in 2-mL tubes containing ceramic beads filled with Tri Reagent (Zymo Research, #R2050-1-200). Tissues were homogenized using an Omni Bead Ruptor 24 and then centrifuged to remove debris. RNA was extracted using the Direct-zol RNA MiniPrep Kit (Zymo Research, #R2052), then converted to complementary DNA (cDNA) using the High-Capacity cDNA Reverse Transcription Kit (Thermo Fisher; #4368813) and RNAse Inhibitor (Takara Bio; #2313B). Total RNA quantity and quality was assessed using the Take3 Microvolume Plate and BioTek Synergy H1 plate reader. Levels of SARS-CoV-2-specific and mouse-specific genes were quantified using PrimeTime™ Gene Expression Master Mix (IDT, #1055771) and TaqMan gene expression primer/probe sets (Applied Biosystems). All qPCR was performed in 384-well plates and run on the ABI QuantStudio 5 real-time PCR system (Applied Biosystems). SARS-CoV-2 RNA-dependent RNA polymerase (RdRp) expression was measured as previously described (Vanderheiden, *et al*., *JVI* 2020). SARS-CoV-2 E gene subgenomic RNA (sgRNA) was measured using a designed forward primer (IDT; 5’-CGATCTCTTGTAGATCTGTTCTC-3’) and the E_Sarbeco R2 reverse primer (IDT; #10006890) and P1 FAM probe (IDT; #10006892).

### Isolation of immune cells from mouse tissues

At day 3 p.i., mice were anesthetized using isoflurane and injected retro-orbitally with CD45:PE (100 µL per mouse, diluted 1:20 in sterile saline; BD Biosciences, clone 30-F11) for intravital labeling of circulating CD45^+^ cells (Anderson, *et al., Nature Protocols* 2014). Five minutes after injection mice were euthanized via isoflurane overdose. The spleen and two lobes of lung tissue were collected from each mouse and placed on ice in RPMI 1640 medium (Corning, #10-040-CM) containing 10% FBS (R10 medium). Spleens were mechanically homogenized, passed through a 70-micrometer (µM) cell strainer (Corning, #431751), and spun down. Cells were treated with ACK lysis buffer (Quality Biological, #118-156-721) for four minutes, washed with R10 medium, and stored on ice for flow cytometry staining. Lungs were mechanically disrupted in six-well culture plates, then digested for 30 minutes at 37°C in a solution of DNase I (final concentration 40 U/mL; Sigma-Aldrich, #DN25) and collagenase (final concentration 100 µg/mL; Sigma-Aldrich, #11088882001) in HBSS. Digestion was stopped with R10 medium, and lungs were pushed through a 70-µM cell strainer to obtain a single-cell suspension. Cells were resuspended in 30% Percoll (Cytiva, #17089101) in DPBS and pelleted. The top layer of cell debris was removed, and the cell pellet was treated with ACK lysis buffer for four minutes. Cells were washed and resuspended in R10 medium and stored on ice for flow cytometry staining.

### Meso Scale Discovery (MSD) assays

Meso Scale Discovery assays were carried out using the V-PLEX SARS-CoV-2 Panel 7 (Mouse IgG) Kit (Meso Scale Discovery, #K15484U-2) per manufacturer protocol. Briefly, 10-spot MSD assay plates were blocked with MSD Blocker A, then washed with 1X MSD Wash Buffer. Serum samples were diluted 1:10,000,000 and added to the plates and incubated for two hours at room temperature with shaking. Following a series of washes, detection antibody was added to the plate and incubated for one hour at room temperature with shaking. The plates were washed with 1X MSD Wash Buffer, and then MSD Gold Read Buffer B was added to each well. Plates were read using an MSD plate reader and analyzed using Discovery Workbench® software, version 4.0.

### Focus Reduction Neutralization Tests (FRNTs)

Mouse sera were collected by submandibular vein bleed just prior to challenge, and FRNT assays were performed as previously described (Vanderheiden, *et al*., *Curr Prot Immunol* 2020). Briefly, serum samples were initially diluted 1:10 and then serially diluted threefold in serum-free DMEM in duplicate. Diluted sera were incubated with an equal volume of MA-SARS-CoV-2 or B.1.351 (with a target of 100 to 200 foci per well) for one hour at 37° C and 5% CO_2_ in a V-bottom 96-well plate. The serum/virus mixture was then added to Vero E6-TMPRSS2 cells cultured in 96-well blackout plates and adsorbed at 37°C and 5% CO_2_ for one hour. After adsorption, the inoculum was removed and replaced with 100 µL per well of pre-warmed 0.85% methylcellulose (Sigma-Aldrich, #M0512-500G) in 1X DMEM. Plates were incubated at 37°C and 5% CO_2_ for approximately 18 hours, after which the methylcellulose overlay was removed and cells washed six times with 1X PBS (prepared from 10X concentrate; Biolegend, #926201). Cells were fixed with 2% PFA in DPBS for 30 minutes. Following fixation, plates were washed twice with 1X PBS. Fixed cells were then incubated with primary antibody directed against the SARS-CoV-2 spike conjugated to AlexaFluor-647 (CR3022-AF647) in permeabilization buffer (0.1% bovine serum albumin, BSA; Fisher, #BP1600-100), and 0.1% saponin (*73-250G-F*), in DPBS at 4°C overnight. Cells were washed three times with 1X PBS, and foci were visualized using the ImmunoSpot CTL analyzer. Antibody neutralization was quantified by counting the number of foci for each sample using the Viridot package in R. The neutralization titers were calculated as 1 - (ratio of the mean number of foci in the presence of sera and foci at the highest dilution of the respective sera sample).

### Histopathological Analysis

Perfused lung sections preserved in 10% neutral buffered formalin were shipped to Histowiz, Inc., where they were paraffin-embedded, sectioned, and mounted onto slides. Slides were stained with H&E or subjected to IHC with an antibody specific for SARS-CoV-2 N protein. Stained slides were read by a board-certified veterinary pathologist at Histowiz (blinded to treatment groups) and scored from 0 to 5 for perivascular inflammation, bronchial/bronchiolar alveolar degeneration/necrosis, bronchial/bronchiolar inflammation, and alveolar inflammation or from 0 to 3 for extent of IHC positivity in the bronchi and the alveoli. A narrative description of leukocyte classes and histopathological findings observed was also provided.

### Flow cytometry analysis

Single-cell suspensions were spun down and resuspended in DPBS with 1% FBS (FACS buffer). Cells were resuspended in FACS buffer containing anti-mouse CD16/32 (Biolegend, clone 93) to block Fc receptors and incubated at 4°C for 15 minutes. A mix of the following antibodies (as well as Ghost Dye 780 live/dead stain, Tonbo Biosciences, #13-0865-T100) diluted in FACS buffer was added, and cells were surface stained at 4°C for 20 minutes: anti-F4/80:FITC (Biolegend, clone BM8), anti-CD64:PerCP-Cy5.5 (Biolegend, clone X54-5/7.1), anti-CD49b:AF647 (Biolegend, clone DX5), anti-I-A/I-E:AF700 (Biolegend, clone M5/114.15.2), anti-CD3ε:APC-Cy7 (Biolegend, clone 145-2C11), anti-CD19:APC-Cy7 (BD Biosciences, clone 1D3), anti-Siglec-F:BV421 (Biolegend, clone S17007L), anti-CD107a:PE-Cy7 (Biolegend, clone 1D4B), anti-CD45.2:BV605 (BD Biosciences, clone 104), anti-Ly6G:BV650 (Biolegend, clone 1A8), anti-CD115:BV711 (Biolegend, clone AFS98), anti-Ly6C:BV785 (Biolegend, clone HK1.4), anti-CD11b:BUV395 (BD Biosciences, clone M1/70), anti-Siglec-H:BUV496 (BD Biosciences, clone 551), anti-CD11c:BUV737 (BD Biosciences, clone HL3), anti-CD86:PE-Dazzle 594 (Biolegend, clone GL-1), anti-NKp46:PE-Cy5 (Biolegend, clone 29A1.4) and anti-H-2Kb:BUV805 (BD Biosciences, clone AF6-88.5). Cells were washed once in FACS buffer and then fixed in 2% PFA in DPBS for 30 minutes at room temperature. Cells were pelleted and resuspended in FACS buffer. Precision Count Beads (Biolegend, #424902) were added to samples to obtain counts, and samples were acquired on a BD FACSymphony A5 flow cytometer. Analysis was performed in FlowJo v10.

### Single-cell RNA-sequencing

Lungs from mice at day 3 p.i. were processed to single-cell suspensions as described above. Single-cell suspensions were subjected to magnetic sorting using MojoSort™ Mouse anti-PE Nanobeads (BioLegend, #480080) to remove IV-positive, PE-tagged cells. Suspensions were subjected to magnetic separation a total of four times before the negative fraction was resuspended in buffer and counted. Cell suspensions were captured in emulsion droplets using the Chromium Next GEM Single Cell 5’ GEM and Gel Bead Kits (10x Genomics, #1000244 and #1000264) on a 10x Chromium Controller. Amplification of cDNA and 5’ GEX library preparation and indexing were performed according to the manufacturer’s instructions using the Library Construction Kit and Dual Index Kit TT Set A (10x Genomics, #1000190 and #1000215). Gene expression libraries were sequenced as paired-end 26-by-91 reads on an Illumina NovaSeq 6000 S4 System targeting a depth of 50,000 reads per cell at the Emory National Primate Research Center Genomics Core. Cell Ranger (v7.0.0) was used for demultiplexing, aligning barcodes, mapping to a custom mm10 reference genome with the SARS-CoV-2 complete genome sequence included (GenBank Accession: MT246667.1), and quantifying unique molecular identifiers. Filtered Cell Ranger matrices were analyzed using the R package Seurat (v4.1). Data were filtered to remove cells with less than 200 expressing genes and greater than 2500 expressing genes. Principal-component analysis and dimensional reduction were conducted on log-normalized and scaled gene expression data. Clustering was conducted using the FindNeighbors and FindClusters functions. Marker genes from Adamska *et al*. were used to identify cell type clusters in the dataset (*35*). The data were downsampled based on the smallest group, resulting in 3,867 cells in each group: mock, naïve infected, and 0.5 μg S-2P-vaccinated infected. Differential gene expression analysis for eosinophils and monocyte/interstitial macrophages was performed using the FindMarkers function in Seurat and top genes were filtered based on adjusted p-val < 0.001 and average log2 fold change > 1. Gene set enrichment analysis (GSEA) was performed with the R package fgsea (v1.20.0) with ranking based on average log2 fold change and 10,000 permutations. Gene expression data using for GSEA were generated using the Seurat FindMarkers function and genelists were obtained from mouse MsigDB (MH, M2, and M5 gene sets) (*36, 74*). Granulocyte cluster markers were determined using the FindAllMarkers function in Seurat using default settings and top markers were filtered based on average log2 fold change. Eosinophil modules were generated using AddModuleScore function in Seurat. Modules were composed of the top 50 genes ranked by log2 fold change from gene lists published in Dolitzky *et al* (*41*).

## Data Availability

Single cell RNA sequencing data will be made publicly accessible through the Gene Expression Omnibus upon publication.

## Quantification and Statistical Analysis

All experiments in mice were repeated twice, with representative results from one experiment shown. Statistical analysis was performed in GraphPad Prism 9.2.0 using Student’s *t* test to compare two groups and unpaired one-way analysis of variance (ANOVA) to compare more than two groups. Statistical analysis for scRNA-seq calculations was performed in R as described above. Throughout the manuscript, results were considered significant below a P value of 0.05.

Antibody neutralization was quantified using the Viridot program to count the number of foci present in each well (Katzelnick, *et al.*, *PLoS Negl Trop Dis* 2018). The neutralization titers were calculated as follows: 1 - (ratio of the mean number of foci in the presence of sera and foci at the highest dilution of respective sera sample). Each specimen was tested in duplicate. FRNT_50_ titers were interpolated using a four-parameter nonlinear regression in GraphPad Prism 9.2.0. Samples that did not neutralize at the limit of detection at 50% were plotted at the LOD and were used for geometric mean and fold-change calculations. RT-qPCR results are expressed relative to mouse *Gapdh* expression for the same sample and were calculated using the ΔΔCT relative quantitation method as compared to mock.

