## Supplementary Figures for "Eosinophils protect against SARS-CoV-2 following a vaccine breakthrough infection"

**Supplemental Figures**

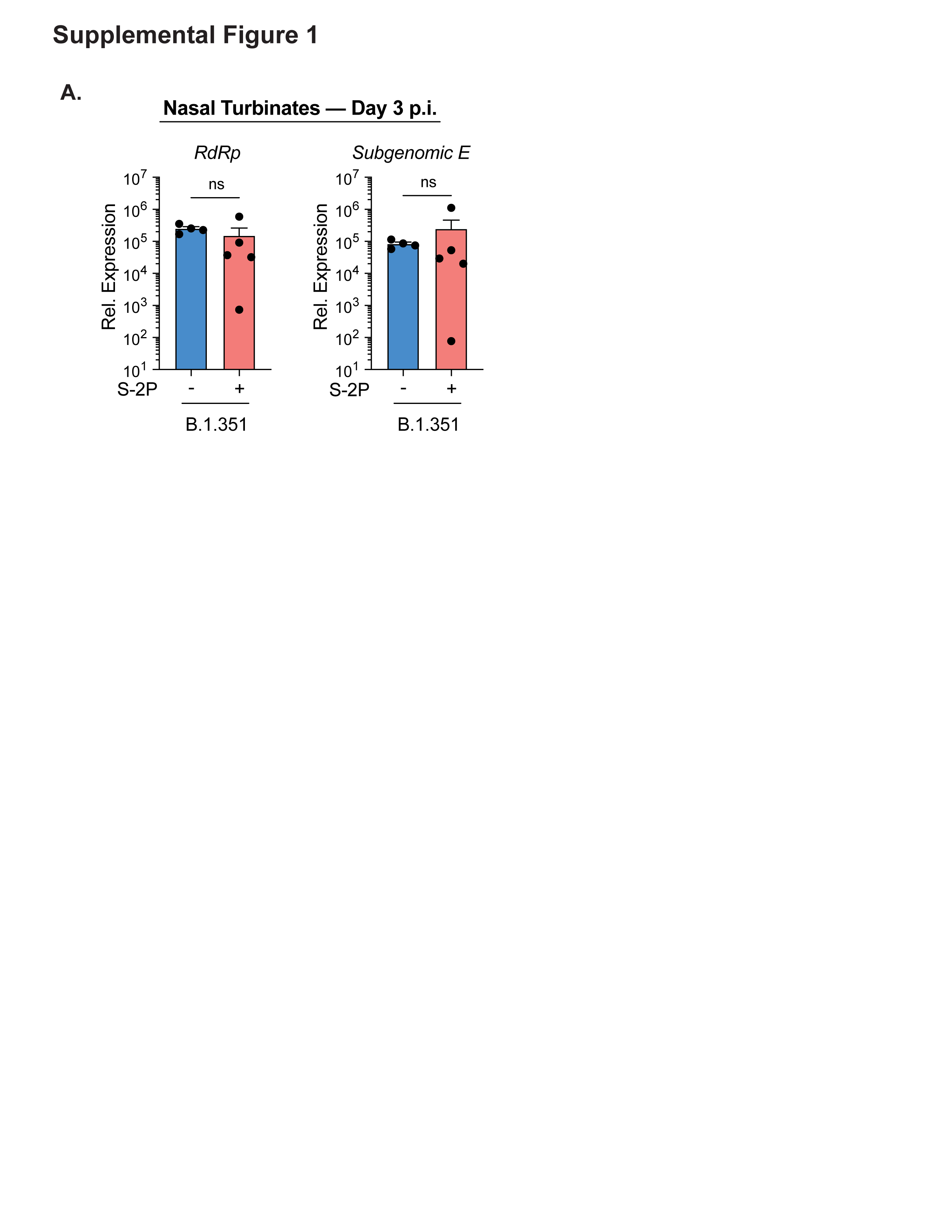
**Supplemental Figure 1: Similar viral load in the nasal turbinates between naïve and low-dose S-2P-vaccinated mice.** B.1.351 viral loads in the nasal turbinates at day 3 p.i. as measured by RdRp genomic RNA by RT-qPCR (left) and sG E gene RNA by RT-qPCR (right). Group color are as follows: naïve infected = blue, 0.5 µg S-2P vaccinated infected = red. Data are represented by the mean +/- the standard error of the mean. Statistical significance was determined using an unpaired Student’s *t* test.

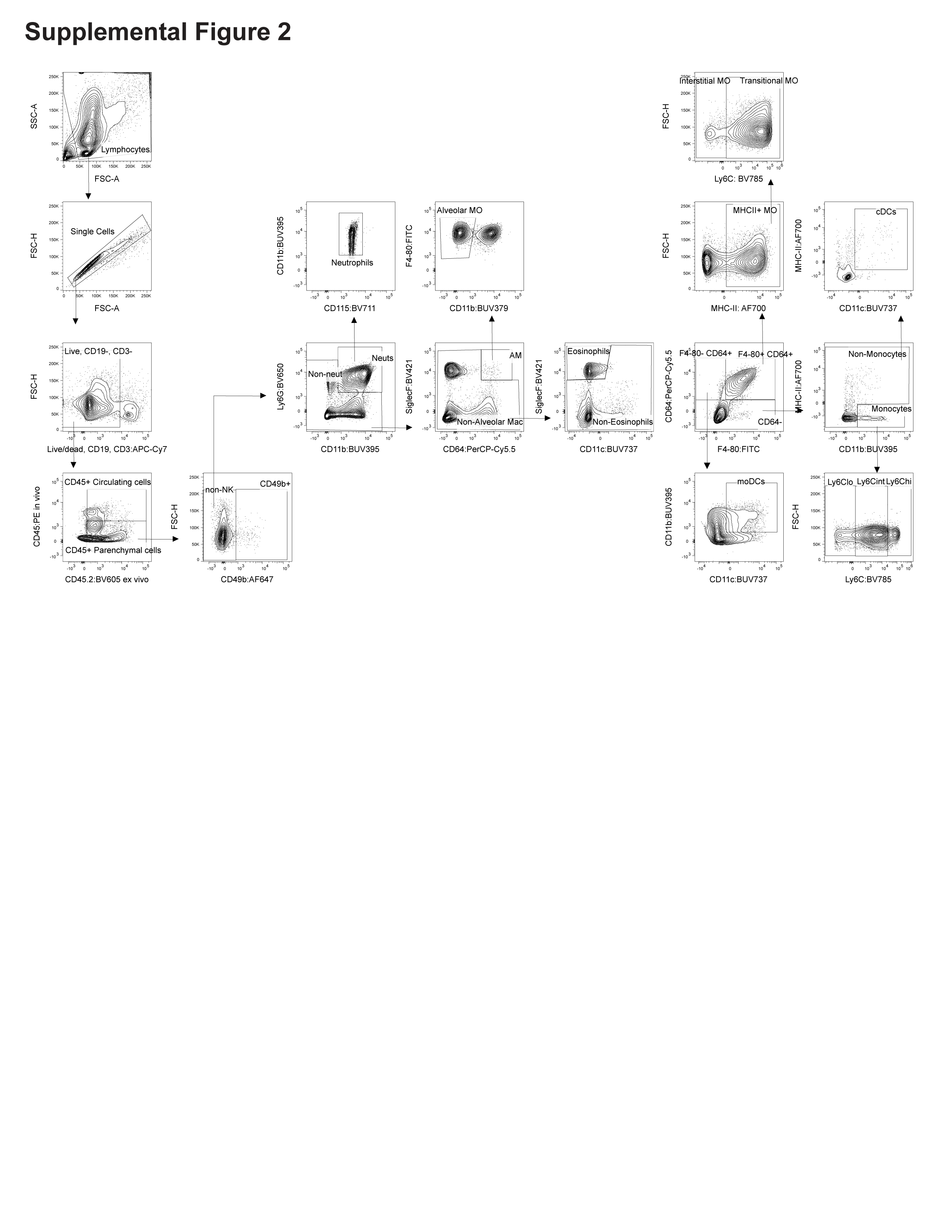
**Supplemental Figure 2: Representative flow cytometry gating strategy of innate immune cell populations in the lungs.** Mice infected with SARS-CoV-2 B.1.351 variant, and lung tissue was harvested at 3 days p.i. Five minutes prior to euthanasia, CD45:PE was injected into mice via the retro-orbital route. Lungs were processed to a single-cell suspension and analyzed via flow cytometry. Cell populations were identified by sequential gating following the black arrows.

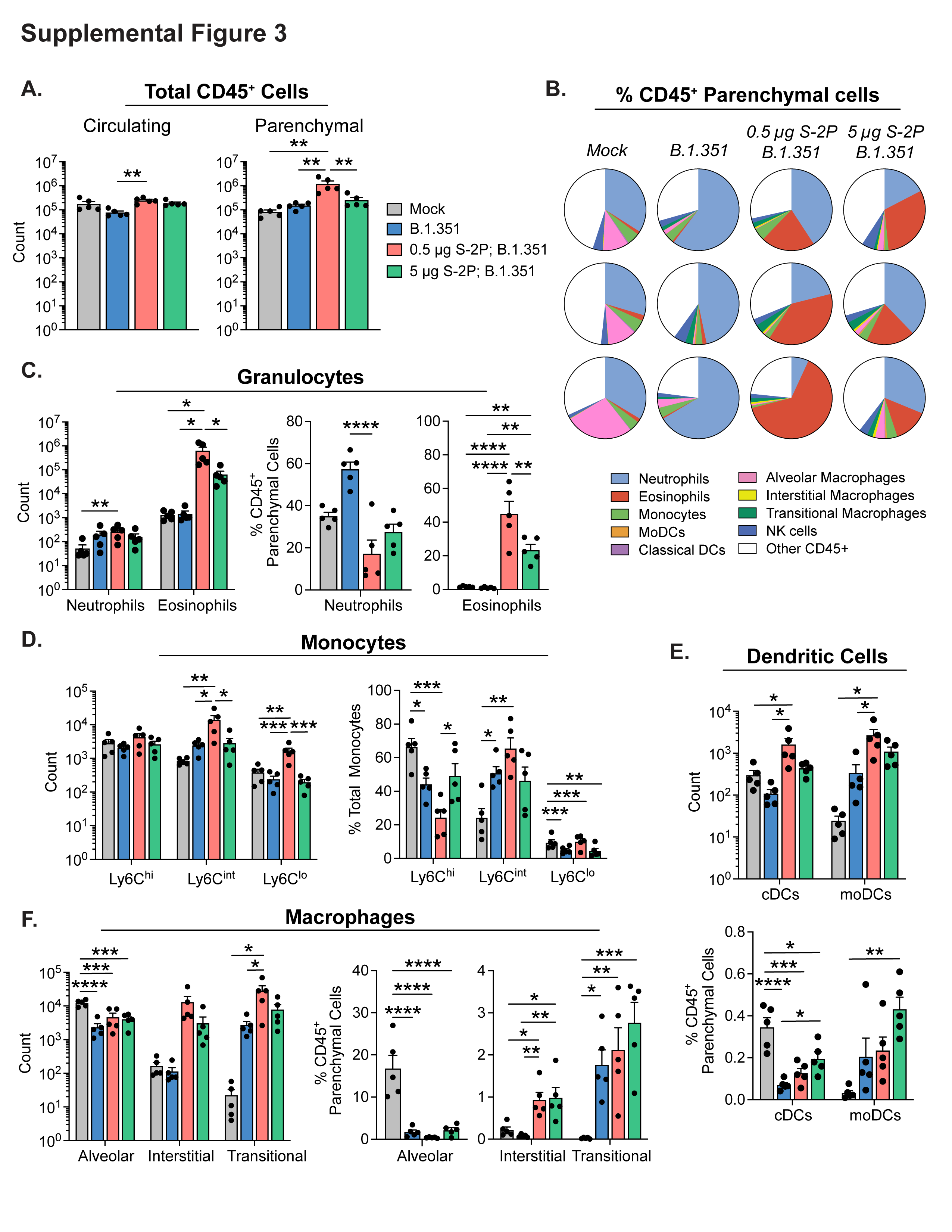

**Supplemental Figure 3. High-dose S-2P vaccination reduces the influx of eosinophils observed in low-dose S-2P-vaccinated mice after B.1.351 infection.** S-2P vaccinated or naïve (unvaccinated) mice were challenged intranasally with 1 x 10^6^ PFU B.1.351 and parenchymal lung immune cells evaluated by flow cytometry at day 3 p.i. (**A**) Absolute counts of CD45^+^ circulating cells (left) and absolute counts of CD45^+^ lung-resident parenchymal cells (right). (**B**) Pie charts displaying proportion of each identified cell type represented as a percent of the total CD45^+^ lung parenchymal cells for 3 representative mice from each group. (**C**) Quantification of granulocytes identified as neutrophils and eosinophils represented as absolute counts (left) and proportion of total CD45^+^ parenchymal cells (right). (**D**) Quantification of monocyte subtypes based on Ly6C expression represented as absolute counts (left) and proportion of total monocytes (right). (**E**) Quantification of classical dendritic cells and monocyte-derived dendritic cells represented as absolute counts (top) and proportion of total CD45^+^ parenchymal cells (bottom). (**F**) Quantification of macrophage subtypes identified as alveolar, interstitial, and transitional macrophages represented as absolute counts (left) and proportion of total CD45^+^ parenchymal cells (right). Group color are as follows: (uninfected) mock = grey, (naïve infected) B.1.351 = blue, (low-dose vaccinated infected) 0.5 µg S-2P, B.1.351 = red, (high-dose vaccinated infected) 5 µg S-2P, B.1.351 = green. Data are represented by the mean +/- the standard error of the mean. Statistical significance was determined using an unpaired one-way ANOVA with Tukey’s multiple comparisons test, and P values are represented above the bar graphs as follows: *, P < 0.05; **, P < 0.01; ***, P < 0.001; ****, P < 0.0001.

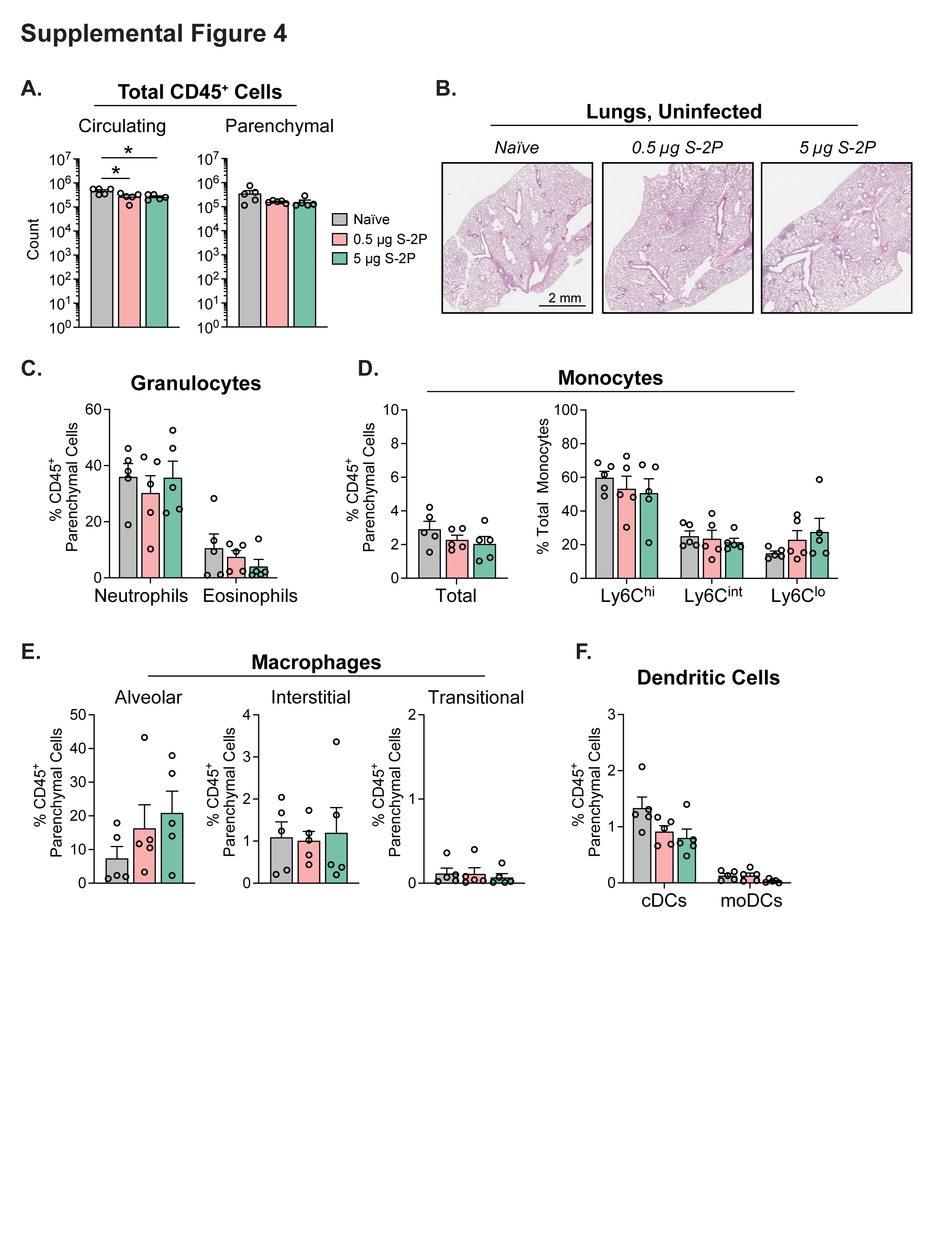
**Supplemental Figure 4. S-2P-vaccinated mice do not exhibit influx of innate immune cells into the lung parenchyma prior to SARS-CoV-2 challenge.** Analysis of lung parenchymal immune cells in uninfected S-2P vaccinated or naïve (unvaccinated) mice. (**A**) Absolute counts of CD45^+^ circulating cells (left) and absolute counts of CD45^+^ lung-resident parenchymal cells (right) determined by flow cytometry at day 3 p.i. (**B**) Histophathologic images of mouse lungs stained with H&E. Scale bars are 2 mm. (**C**) Quantification of granulocytes identified as neutrophils and eosinophils represented as proportion of total CD45^+^ parenchymal cells. (**D**) Quantification of monocyte subtypes based on Ly6C expression represented as proportion of total CD45^+^ parenchymal cells (left) of total monocytes (right). (**E**) Quantification of macrophage subtypes identified as alveolar, interstitial, and transitional macrophages represented as proportion of total CD45^+^ parenchymal cells. (**F**) Quantification of classical dendritic cells and monocyte-derived dendritic cells represented as proportion of total CD45^+^ parenchymal cells. Group color are as follows: (unvaccinated) naïve = grey, (low-dose vaccinated) 0.5 µg S-2P, B.1.351 = light red, (high-dose vaccinated) 5 µg S-2P, B.1.351 = light green. Data are represented by the mean +/- the standard error of the mean. Statistical significance was determined using an unpaired one-way ANOVA with Tukey’s multiple comparisons test, and P values are represented above the bar graphs as follows: *, P < 0.05.

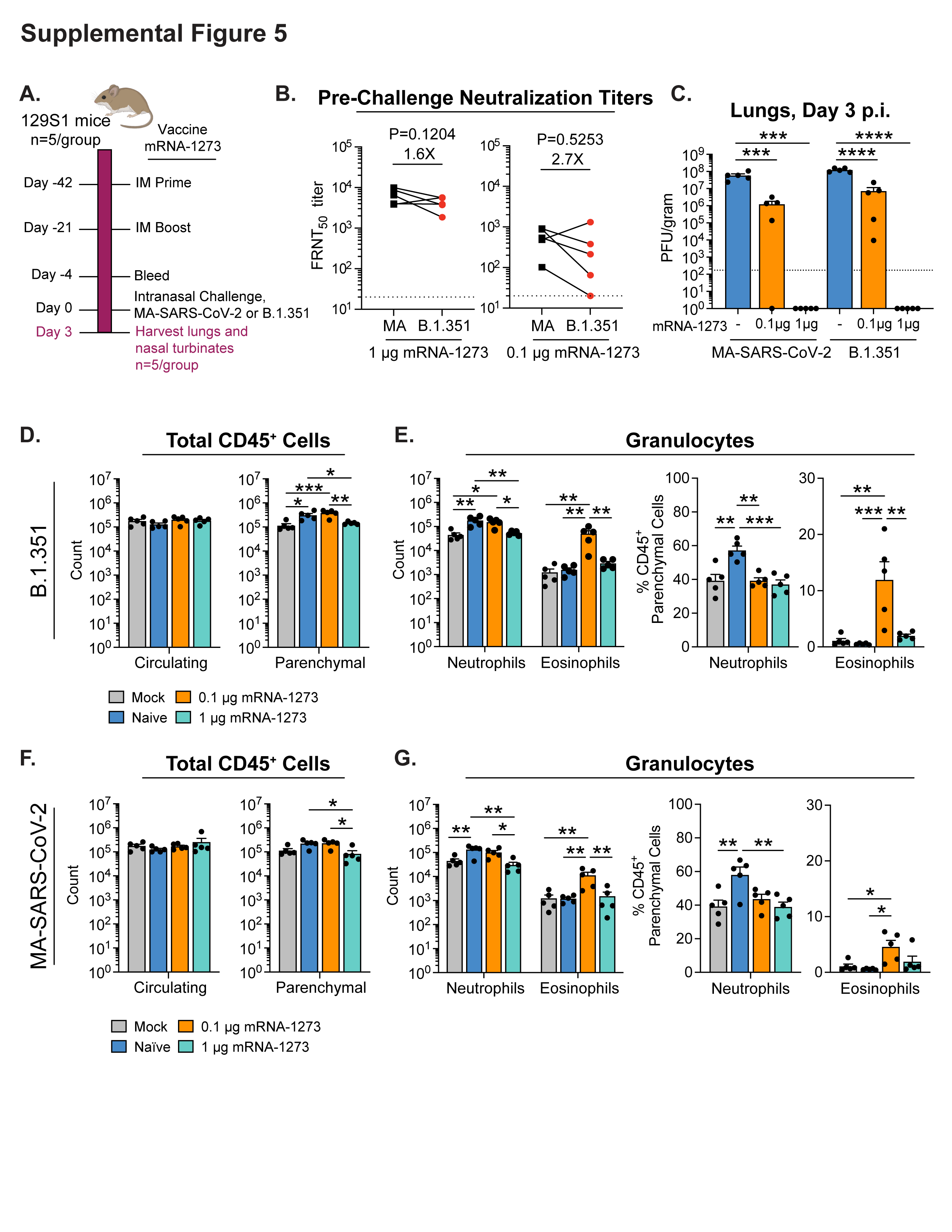
**Supplemental Figure 5. Vaccination with mRNA-1273 prior to SARS-CoV-2 challenge provides viral control and spurs eosinophil infiltration.** Experimental schematic detailing vaccination of 129S1/SvImJ mice with mRNA-1273 and subsequent intranasal challenge with 1 x 10^6^ PFU B.1.351 or mouse-adapted SARS-CoV-2 (MA-SARS-CoV-2). Parenchymal lung immune cells evaluated by flow cytometry at day 3 p.i. (**A**) Experimental schematic detailing vaccination of 129S1/SvImJ mice with low (0.1 µg) and high (1 µg) doses of mRNA-1273 and subsequent challenge with B.1.351. (**B**) Live virus serum FRNT_50_ titers against MA-SARS-CoV-2 (black circles) or B.1.351 (red circles) at 4 days before challenge in mice vaccinated with the indicated dose of mRNA-1273. Connected circles represent the same mouse. The P value and fold difference in GMT between MA-SARS-CoV-2 and B.1.351 for each dose of mRNA-1273 are shown. The dotted horizontal line represents the LOD of the assay. (**C**) Viral loads in the lungs as measured by plaque assay at day 3 p.i. (**D-E**) Quantification of lung immune cells by flow cytometry 3 days after challenge with B.1.351. (**D**) Absolute counts of CD45^+^ circulating cells (left) and absolute counts of CD45^+^ lung-resident parenchymal cells (right). (**E**) Quantification of parenchymal granulocytes identified as neutrophils and eosinophils represented as absolute counts (left) and proportion of total CD45^+^ parenchymal cells (right). (**F-G**) Quantification of lung immune cells by flow cytometry 3 days after challenge with MA-SARS-CoV-2. (**F**) Absolute counts of CD45^+^ circulating cells (left) and absolute counts of CD45^+^ lung-resident parenchymal cells (right). (**G**) Quantification of parenchymal granulocytes identified as neutrophils and eosinophils represented as absolute counts (left) and proportion of total CD45^+^ parenchymal cells (right). Group color are as follows: (uninfected) mock = grey, (unvaccinated infected) naïve = blue, (low-dose vaccinated infected) 0.1 mRNA-1273 = orange, (low-dose vaccinated infected) 1 µg mRNA-1273 = teal. Data are represented by the mean +/- the standard error of the mean. Statistical significance was determined using an unpaired one-way ANOVA with Tukey’s multiple comparisons test, and P values are represented above the bar graphs as follows: *, P < 0.05; **, P < 0.01; ***, P < 0.001. Created with BioRender.com.

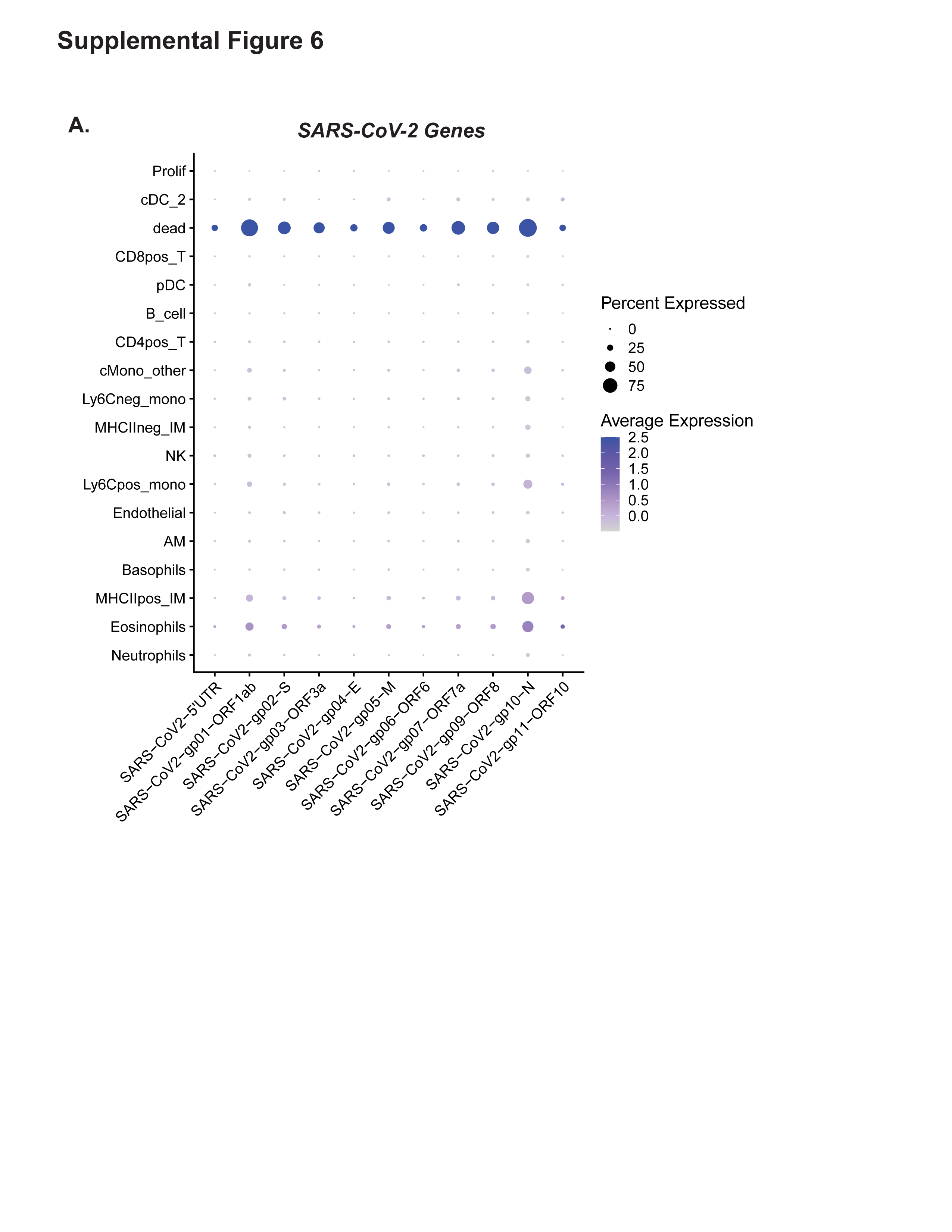
**Supplemental Figure 6. Expression of SARS-CoV-2 genes across samples by scRNA-seq.** At day 3 p.i. lung parenchymal cells were isolated and evaluated by scRNA-seq. The average expression and percent of cells expressing each SARS-CoV-2 gene detected was determined for each cell type. These data are represented as a bubble plot, where the size of the circle represents the percent of cells expressing the given gene and the color of the fill represents the average expression among those cells.

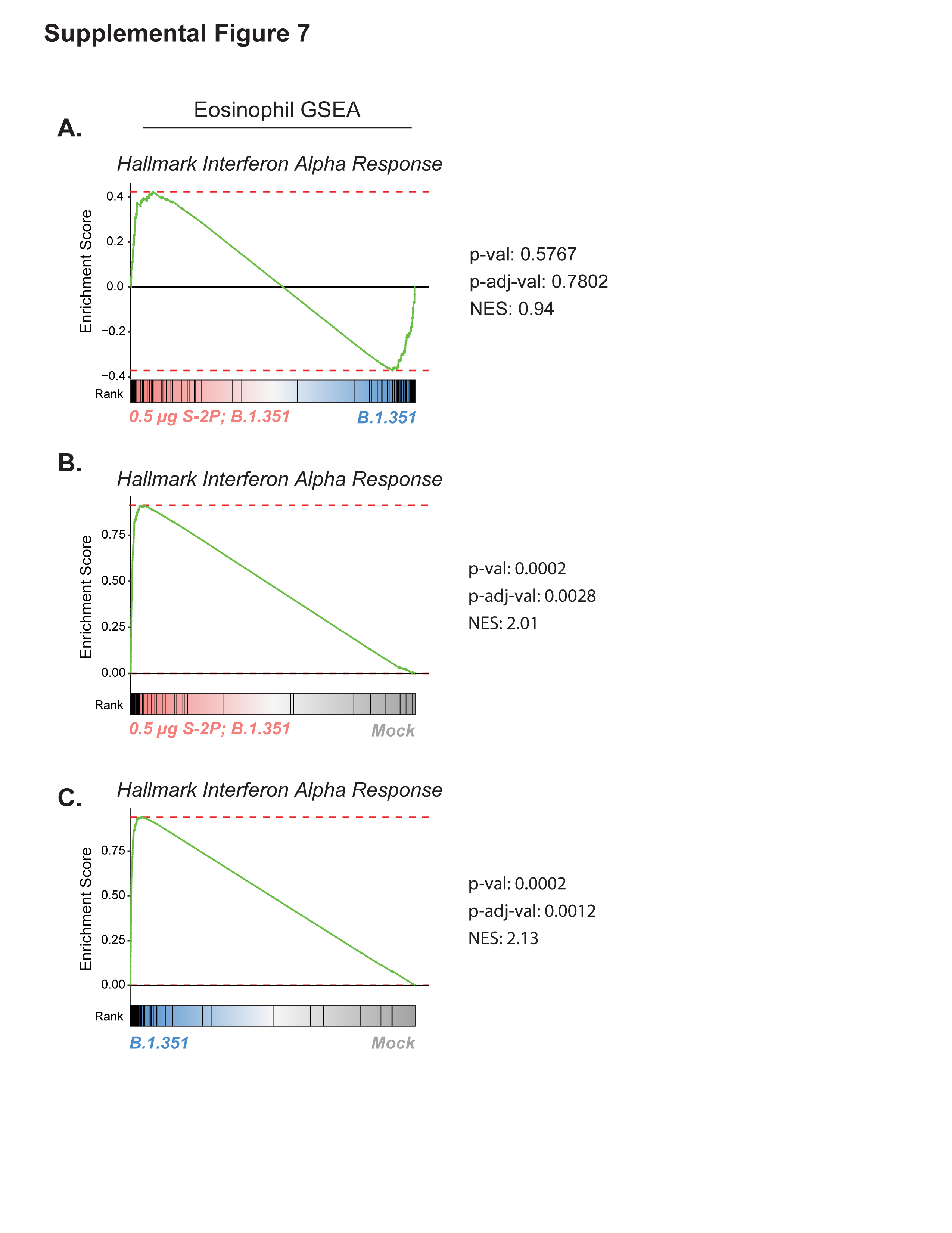
**Supplemental Figure 7. Gene set enrichment analysis (GSEA) for Hallmark Interferon Alpha Response pathway on subsetted granulocytes.** GSEA analysis plots for group pairs: (A) 0.5 μg S-2P-vaccinated infected vs naïve infected, (B) 0.5 μg S-2P-vaccinated infected vs mock, (C) naïve infected vs mock.

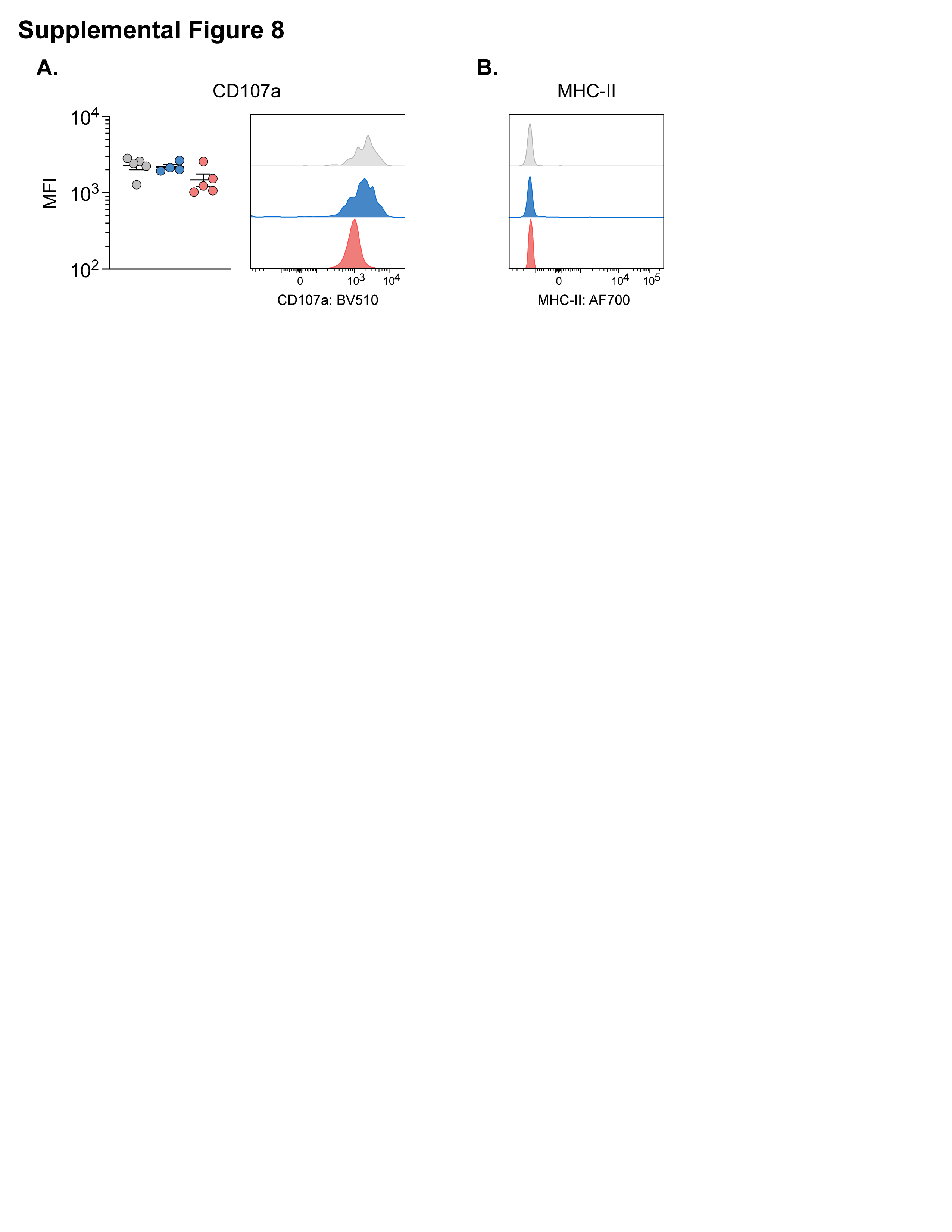
**Supplemental Figure 8. Flow cytometry analysis of lung parenchymal eosinophil cell surface expression of CD107a and MHC-II at 3 days p.i.** Plots displaying mean fluorescence intensity (MFI) (left). Representative histograms of expression for each group (right). Group color are as follows: (uninfected) mock = grey, (naïve infected) B.1.351 = blue, (vaccinated infected) 0.5 µg S-2P, B.1.351 = red.
